# Neuropeptide signalling and perineurial barrier generate a persistent stress-induced internal state in *Drosophila*

**DOI:** 10.1101/2025.05.08.652787

**Authors:** Abdalla G. Alia, Xinyue Hu, Yuzhe Gu, Guangnan Tian, Julie L. Semmelhack, Kokoro Saito, Hiromu Tanimoto, Koki Tsuyuzaki, Shintaro Naganos, Tomoyuki Miyashita, Minoru Saitoe, Yukinori Hirano

## Abstract

Although fear conditioning has elucidated cue-evoked acute fear responses, the mechanisms by which stress experiences induce generalized internal states linked to anxiety are poorly understood. Here, we report that robust stress induces a persistent behavioral change characterized by avoidance of a confined space, claustrophobia-like behavior in *Drosophila*. Unlike aversive memory formation, the development of claustrophobia-like behavior does not require dopamine receptors. Our neuronal screening determined that neuropeptide signalling via Allatostatin-A inactivates the downstream neurons via its receptor AstA-R1, causally inducing claustrophobia-like behavior. Moreover, gene expression profiling of individual fly heads revealed that immune response activation in perineurial barrier is involved in claustrophobia-like behavior. Our data demonstrate that stress-induced persistent behavioral change would not be related to a canonical mechanism of aversive memory formation, rather involves neuropeptidergic signalling and perineurial barrier, providing the mechanism determining internal states which persistently change behavioral modes.

## Introduction

Robust stresses are significant risk factors for the development of psychiatric disorders, including anxiety, depression, and post-traumatic stress disorder (PTSD). Traditionally, fear conditioning, where contextual cues are associated with an aversive stimulus has been extensively employed to investigate mechanisms of fear memory that elicit acute fear responses, such as freezing and avoidance, in various model animals ^1, 2^. In addition to acute fear responses, stress experiences also induce sustained and generalized states ^2^ as seen in various types of phobias ^3^, such as agoraphobia and claustrophobia, which evoke uncontrollable, persistent, and excessive fear in response to potentially harmful contexts or objects. The evolutionary advantage of phobic states would be to enhance risk assessment in life-threatening situations to avoid potential dangers. Understanding the mechanisms of how a stressful event persistently changes the brain states that switch the behavioral modes to the one related to anxiety or phobia is one of the major challenges in alleviating the symptoms of psychiatric disorders.

In rodents, the study of generalized anxiety has expanded beyond fear conditioning to include various behavioral assays, such as the elevated plus maze, open field, and light/dark box, providing insights into stress-induced alterations in exploratory and avoidance behaviors^4^ that can be related to phobic behaviors in humans. Various neuron types in the amygdala^5, 6^, locus coeruleus (LC) ^6, 7^, lateral septum^8^, hypothalamus^9^, and claustrum^10^ have been proposed to be causally linked to stress-induced anxiety, by employing different stress models such as restraint^6, 8^, foot shock^7, 9, 11^, and social defeat stress^10, 12, 13, 14^. These studies highlight the essential roles of multiple signaling mediated by corticotropin-releasing hormone (CRH) ^6^, norepinephrine (NE) ^7^, Galanin^7^, and tachykinin 2^14^, further suggesting that anxiety state is generated by recruiting distributed array of the anxiety-associated neural circuits^4^. Due to this complexity, the core mechanism that initially alters the brain state remains enigmatic. As the traits of anxiety are likely evolutionarily conserved, model organisms with simpler neural and gene networks may be well-suited to screen and uncover the core and conserved components inducing stress-dependent generalized state.

In *Drosophila*, an aversive training paradigm in which olfactory or visual cues are associated with electric shocks has been utilized to study the neuronal and molecular mechanisms of acute avoidance response which involves the central roles of the memory center, mushroom body neurons^15, 16^ and the dopamine receptor Dop1R1^17^. However, the persistent behavioral changes unrelated to the conditioned stimuli after electric shocks have not been explored. Instead, a continuous vibration-stress has been applied to induce depression-like behavior, which was depicted by reduced climbing attempts over a spatial gap ^18^. Another behavior that could assess anxiety is wall-following behavior ^19^. Persistent behavioral changes have been also reported after repeated fighting losses ^20^ and sexual deprivation ^21^. However, the core mechanisms of stress-dependent generalized state has not been revealed in fly models.

Here, we present a quantitative behavioral assay of active choice during exploration in which *Drosophila* exhibits persistent claustrophobia-like behavior after stress exposure. Notably, this behavior does not require dopamine receptor signalling, suggesting a divergence from canonical fear memory pathways. Our neuronal screening showed that claustrophobia-like behavior requires neuropeptide signalling via Allatostatin-A (AstA) and its receptor AstA-R1, a homolog of mammalian galanin receptor ^22^ that is targeted by treatments for anxiety and depression ^23^. Furthermore, immune signalling mediated by Toll receptors in the perineurial barrier, a component of the fly blood-brain barrier, was implicated in the development of this phenotype. Our findings revealed neuronal and perineurial barrier mechanisms causally related to stress-induced internal states.

## Results

### Electric shock exposure induces claustrophobia-like behavioral phenotype

Animals explore their environment while being alert to potential dangers, which can be influenced by their internal state as seen in the rodent study using the elevated plus maze or open field^4^. To examine stress-induced exploratory behavioral changes in flies, we designed a two-arm arena where male flies explore freely within a 15 mm-diameter central zone connecting to two arms with different widths, 4 mm and 2 mm, slightly larger than male body width of 1.5 mm (Fig. 1a). Male flies were exposed to pulsed electric shocks, 1.5 sec-shock at 60 V with 3.5 sec-rest which cycle was continued for 5 min, in a tube of Φ 14 mm x L 80 mm. After they rested for 1.5 h in food vials, their exploration behavior was examined in the two-arm arena. While naive flies explored both arms equally, shocked flies exhibited reduced exploration in the narrower arm (Fig. 1b, c). Relative time spent in the wide arm compared to the narrower arm (performance index, Fig. 1d) indicated that the preference to the wider arm is significantly induced by shock exposure (Fig. 1d). Shock exposure decreased the cumulative duration spent in both arms, with a more pronounced decrease in the narrower arm (Fig. 1e), indicating active avoidance behavior of narrower spaces. Shock exposure also significantly decreased locomotor activity (Fig. 1f), resulting in reduced entries into both arms (Fig. 1g). However, shocked flies exhibited similar entry frequency into both wide and narrow arms (Fig. 1g). Therefore, the preference against the narrower arm after the shock is not due to a general strategic change in exploration. This behavior involves visual spatial perception, because the shocked flies did not exhibit significant preference in the two-arm arena in dark (Supplementary Fig. 1). Given that this behavior may be analogous to claustrophobia in humans ^24^, we refer the preference against the narrower space as claustrophobia-like behavior.

**Fig. 1.**
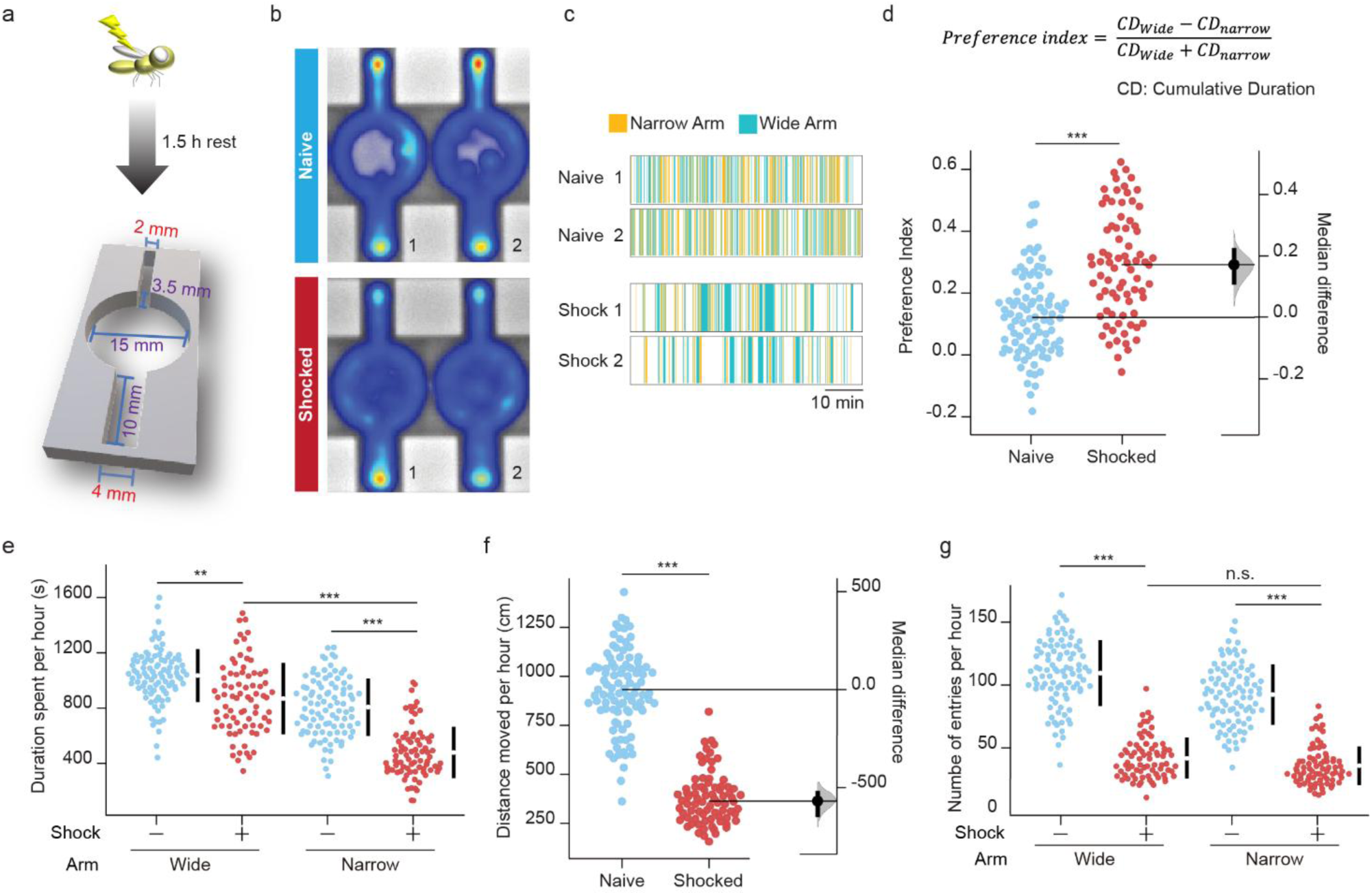
Electric shock exposure induces claustrophobia-like behavior. **a**, Schematic view of an experimental arena. Naïve flies or shocked flies at 60 V for 5 min, which were rested for 1.5 h, were video-recorded for 1 h to analyze the behavior (b-g). **b,c**, Heatmap (b) and time course visualization (c) of representative fly behavior reveal distinct patterns. The time spent in narrow and wide arms in 1 h-recording periods was indicated in orange and green, respectively (c). **d,** Preferential exploration to wide arm is induced by shock exposure. The arm-choice bias of which flies stay in the wide arm was calculated by the indicated formula, resulting in the preference index. Two-sided Mann-Whitney U-test; *n* = 78-95. **e,** Exploration to narrow arm is robustly reduced by shock exposure. Total duration of time spent in each arm in 1 h was presented. Kruskal-Wallis test, *P* <0.0001; *n* = 78-95. **f,** Locomotor activity is reduced by shock exposure. Total distance (cm) moved in the entire arena in 1 h was presented. Two-sided Mann-Whitney U-test; *n* = 78-95. **g,** Entry to narrow arm is not different from that to wide arm after shock exposure. The entry events to both arms were counted in 1 h. Kruskal-Wallis test, *P* <0.0001; *n* = 78-95. Data are represented as mean ± s.e.m., with median differences based on bootstrap estimation. n.s., not significant *P* > 0.05; *, *P* < 0.05; **, *P* < 0.01; ***, *P* < 0.001.

### Claustrophobia-like behavior demonstrates manifestation of anxiety-like internal states, which is not regulated by mechanisms of associative memory formation

We further characterized the effect of aversive stimuli on the development of claustrophobia-like behavior. The intensity and duration of the electric shocks positively correlated with the degree of claustrophobia-like behavior (Fig. 2a, 2b), demonstrating that the development of this behavior is scalable with the aversiveness of the experience. Similarly, heat shock at 40°C for 10 min induced claustrophobia-like behavior (Fig. 2c), but other stressors including vibration ^18^ (Supplementary Fig. 2a) and restraint ^25^ (Supplementary Fig. 2b) did not. The variability of the claustrophobia-like behavior within the population (Fig. 1d) may be due to individual differences or a randomness of phenotypic emergence. Reanalysis of each 20-min time window indicated that individual flies exhibited persistent specific degrees of claustrophobia-like behavior throughout the testing period (Fig. 2d), demonstrating that the variability is due to persistent individuality over time. These characteristics in scalability, generalizability, and persistence suggest that claustrophobia-like behavior includes properties to infer stress-induced internal states ^26^ which may reflect anxiety-like state.

**Fig. 2.**
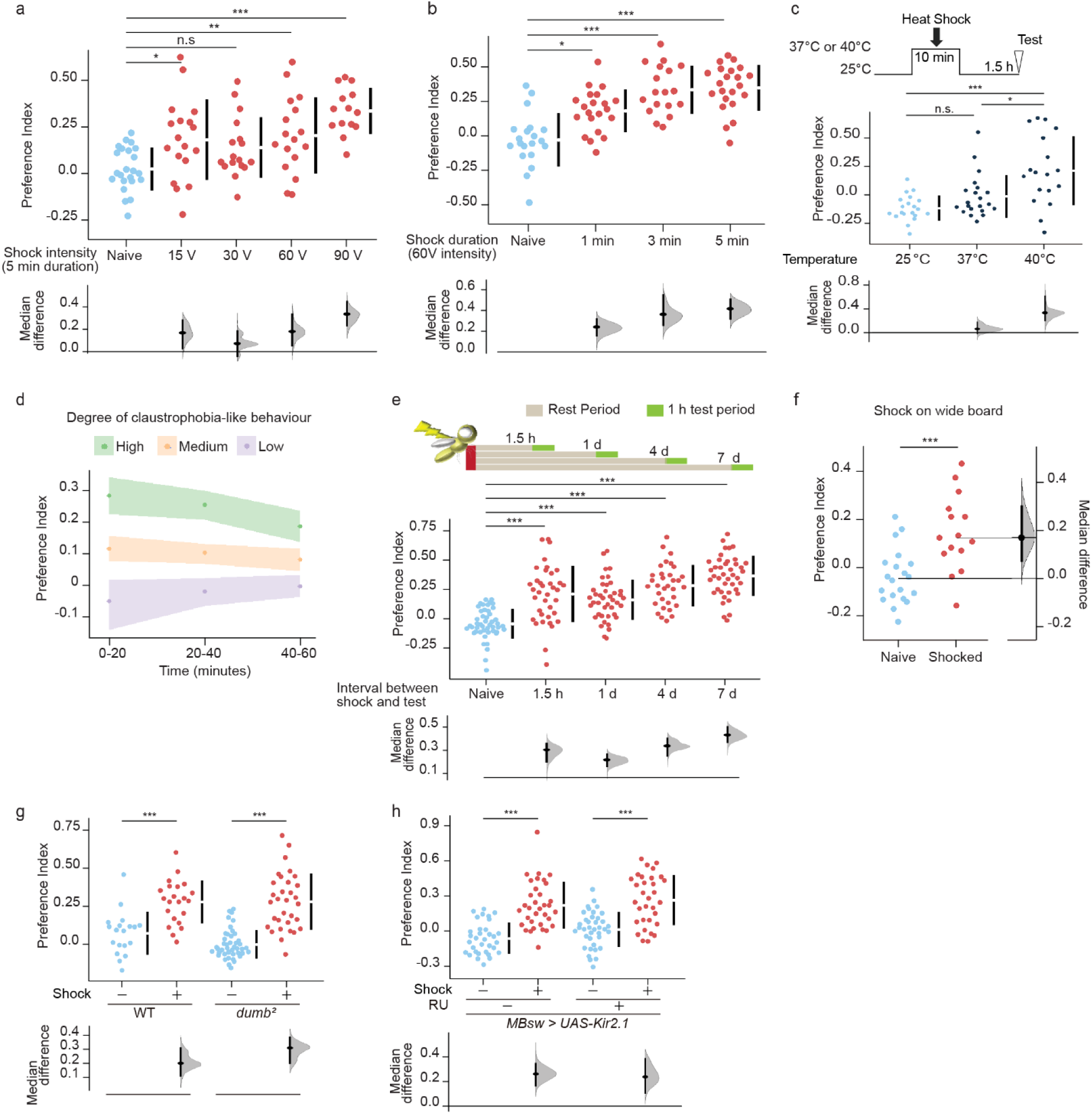
Claustrophobia-like behavior is induced according to the degree of aversive experiences, which does not involve mechanisms of associative memory formation. **a-c**, Development of claustrophobia-like behaivor correlates to the robustness of aversive stimuli. Flies were subjected to electric shocks for various intensities (a) or different durations (b), or heat shocks (c) and rested for 1.5 h. Exploration behavior was analyzed in the two-arm arena. Kruskal-Wallis test, a, *P* < 0.0001, *n* = 14-18; b, *P* < 0.0001, *n* = 18-22; c, *P* = 0.0003, *n* = 18-21. **d,** Individual flies persistently show similar degree of claustrophobia-like behavior. The flies analyzed in Fig. 1d were grouped according to the preference indices with a cutoff below 0.05 as low, that above 0.15 as high, and the rest as medium. Each group was analyzed by splitting into each 20-min window. **e,** The claustrophobia-like behavior persists for 7 days. Flies exposed to electric shocks at 60 V for 5 min were rested for the indicated durations of time before testing their behavior in the two-arm arena. Kruskal-Wallis test, a, *P* < 0.0001; *n* = 33-47. **f,** Flies shocked on a wide electric grid exhibited claustrophobia-like behaviour. Flies were subjected to electric shocks in a space with 120 mm L x 86 mm W x 3.5 mm H at 60 V for 5 min. The flies rested for 1.5 h were analysed in the two-arm arena. Two-sided Mann-Whitney U-test; *n* = 20, 15. **g,h,** Mutation in *Dop1R1* or suppression of MB neural activity does not affect the development of claustrophobia-like behavior. The indicated flies were shocked at 60 V for 5 min, rested for 1.5 h, and tested in the two-arm arena. RU486 (RU) was fed for 2 days before shock exposure. Kruskal-Wallis test, a, *P* < 0.0001; *n* =32-38. Data are represented as mean ± s.e.m., with median differences based on bootstrap estimation. n.s., not significant *P* > 0.05; *, *P* < 0.05; **, *P* < 0.01; ***, *P* < 0.001.

Whereas aversive olfactory associative memory produced by continuous learning for 5 min, known as massed training, does not last for more than 4 days ^27^, the claustrophobia-like behavior induced by electric shocks for 5 min persisted for 7 days post-shock (Fig. 2e). This suggests that the mechanisms inducing claustrophobia-like behavior might differ from those required for aversive associative memory formation. Nonetheless, the flies could associate the shock with the confined space of the tube (14 mm Φ x 80 mm L), resulting in the avoidance from the 2 mm arm. However, the flies shocked in the larger context by using a relatively broad electric grid (120 mm L x 86 mm W x 3.5 mm H) still exhibited claustrophobia-like behavior (Fig. 2f), suggesting that associative memory is not required for the claustrophobia-like behavior. Dopamine receptor 1 (Dop1R1) is essential for associative aversive memory formation ^17^. Importantly, *dumb*^2^ mutant flies which do not show Dop1R1 expression^17^ and observable associative memory, significantly developed the claustrophobia-like behavior after shock (Fig. 2g), suggesting that Dop1R1 receptor is dispensable for developing claustrophobia-like behavior. Additionally, silencing mushroom body neurons required for encoding aversive associative memory in flies^15, 16^, by expressing the human inwardly rectifying K^+^ channel Kir2.1^28^ under the control of mushroom body gene-switch (MBsw) ^29^, which is activated only upon administration of the pharmacological ligand RU486, did not affect the shock-induced claustrophobia-like behavior (Fig. 2h). Taken together, these observations suggest that claustrophobia-like behavior is not a result of memories associating confined spaces with aversive experiences, but rather an anxiety-like internal state.

### A pair of Allatostatin-A neuropeptidergic neurons responds to electric shock and induces the development of claustropbohia-like behavior

To investigate the neuronal mechanisms developing claustrophobia-like behavior, we targeted neuropeptidergic neurons, which could regulate a wide range of behavioral states ^30^. Activity of different populations of neuropeptidergic neurons was suppressed by expressing Kir2.1 under the control of the 14 available neuropeptide GAL4 drivers ^31^. Claustrophobia-like behavior was significantly suppressed by silencing the neurons expressing Allatostatin A (AstA) (Fig. 3a, Supplementary Fig. 3a). Similar suppression was observed by silencing the neurons expressing Myosuppressin (Ms), Corazonin (Crz), and Drosulfakinin (Dsk) to a lesser extent (Fig. 3a, Supplementary Fig. 3a). Additionally, silencing AstA partially alleviated the reduction in locomotor activity (Supplementary Fig. 3b). Given that the suppression of AstA-releasing neurons showed the most significant effect on claustrophobia-like behavior and that AstA is related to mammalian galanin ^22^, which has been linked to anxiety and depression ^23^, we subsequently focused on the role of AstA signaling.

**Fig. 3.**
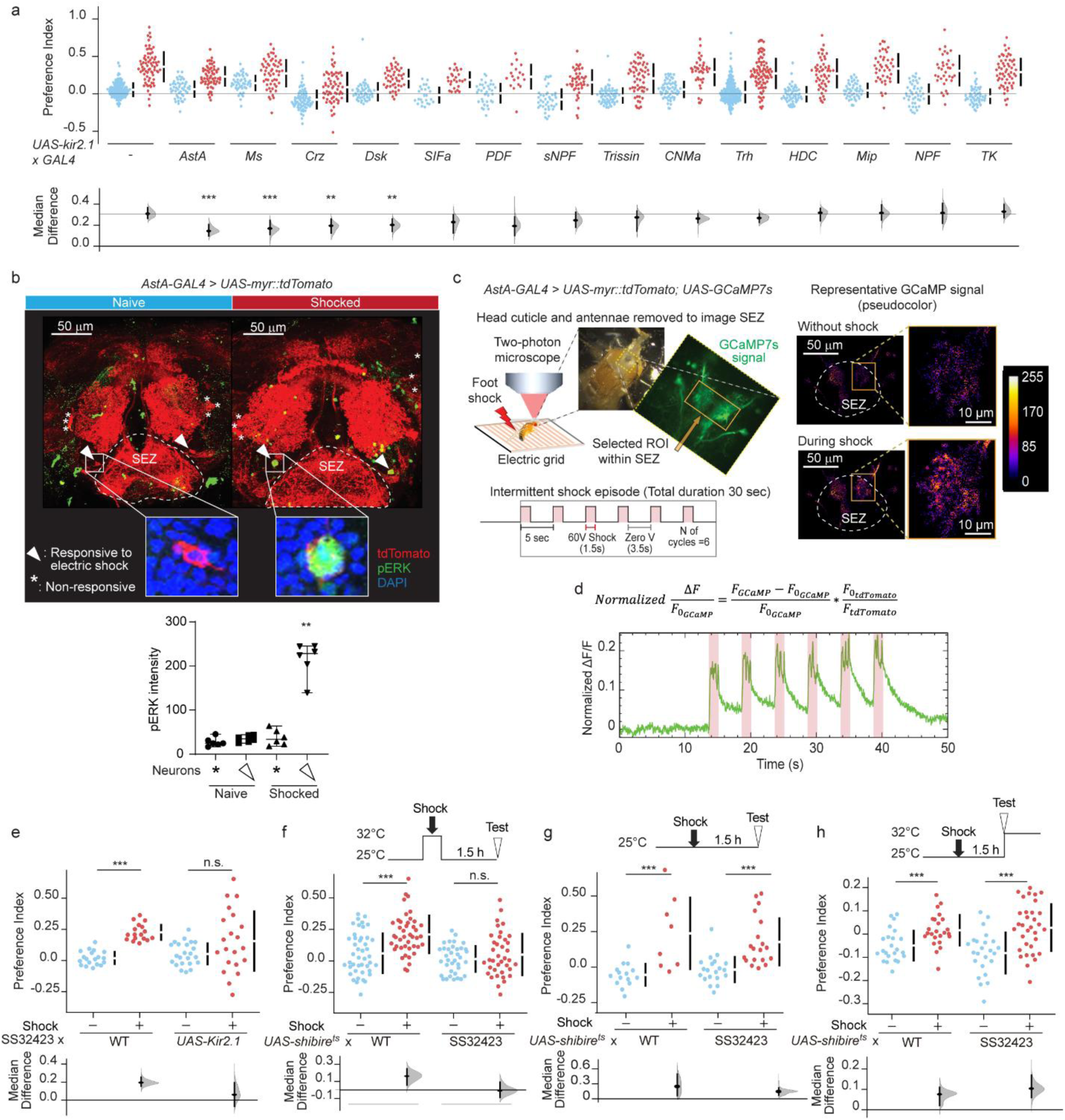
A pair of Allatostatin-A neuropeptidergic neurons is required for stress-induced claustrophobia-like behavior. **a**, Screening for the neuropeptidergic neurons responsible for claustrophobia-like behavior. The indicated flies were subjected to electric shocks at 60 V for 5 min, rested for 1.5 h, and tested in the two-arm arena. Kruskal-Wallis test, a, *P* < 0.0001; *n* = 18-155. **b,** Nuclear pERK is induced by electric shocks. The indicated flies were shocked at 60 V for 5 min. The brains were fixed 10 min after the shock and stained with anti-pERK antibody. Kruskal-Wallis test, a, *P* = 0.0026; *n* = 6 for all. **c,** Schematic of two-photon calcium imaging. Head cuticle and antennae were removed to image the SEZ region. Six cycles of 1.5-sec shocks with 3.5-sec rest intervals were delivered to the flies. The images are representative of two experimental replicates. **d,** Normalized GCaMP7s signals. The GCaMP7s intensities were normalized by the tdTomato intensities, which was subtracted by the normalized GCaMP7s values before shock exposure. The trace is representative of two experimental replicates. **e-h,** Inactivating neurons labelled by SS32423 impaired the development of claustrophobia like behavior. The indicated flies were subjected to electric shocks at 60 V for 5 min, rested for 1.5 h, and tested in the two-arm arena. The temperature was shifted as indicated. Kruskal-Wallis test, e, *P* < 0.0001, *n* = 17-23; f, *P* < 0.0001, *n* = 36-44; g, *P* < 0.0001, *n* = 8-20; h, *P* < 0.0001, *n* = 23-33. Data are represented as mean ± s.e.m., with median differences based on bootstrap estimation. n.s., not significant *P* > 0.05; *, *P* < 0.05; **, *P* < 0.01; ***, *P* < 0.001.

We next ask whether the AstA-ergic neurons are activated by shock, using the phosphorylation of extracellular signal-related kinase (pERK) in the nucleus as a readout ^32^. One pair of *AstA-GAL4*-labelled neurons innervating the suboesophageal zone (SEZ) showed robust pERK signals observed 10 min after shock exposure (Fig. 3b), hereafter referred to as electric shock (ES)-responsive AstA-ergic neurons. We further monitored the activity of *AstA-GAL4*-labelled neurons in the SEZ by expressing the calcium indicator, GCaMP7s ^33^, along with red fluorescent protein myr::tdTomato (Fig. 3c). The GCaMP7s signals were robustly increased in the neurites in the SEZ in response to electric shocks (Fig. 3d), demonstrating that *AstA-GAL4*-labelled neurons are activated by electric shocks.

To specifically manipulate ES-responsive AstA-ergic neurons, we found that the split GAL4 driver SS32423 ^34^ labelled the ES-responsive AstA-ergic neurons along with a few off-target non-responsive neurons (Supplementary Fig. 3c). Silencing SS32423 neurons via expressing Kir2.1 impaired the development of claustrophobia-like behavior (Fig. 3e, Supplementary Fig. 3d) without rescuing the decreased locomotor activity compared to controls (Supplementary Fig. 3e), demonstrating a causal role of these neurons in the development of claustrophobia-like behavior. To temporally suppress the synaptic output, *shibire^ts^* ^35^, a temperature-sensitive variant of dynamin was expressed by SS32423, which blocks synaptic transmission at temperatures of 29°C or higher. Applying electric shock at 32°C significantly reduced the claustrophobia-like behavior (Fig. 3f, Supplementary Fig. 3f), while control experiments under continuous 25°C or treatment at 32°C during test did not affect the claustrophobia-like behavior (Fig. 3g, 3h). Therefore, activity of ES-responsive AstA-ergic neurons during shock exposure is crucial for the development of claustrophobia-like behavior. Moreover, Kir2.1-mediated silencing of SS32423 did not affect the performance of flies in olfactory aversive memory test (Supplementary Fig. 3g), suggesting a dissociation between memory functionality and stress-induced brain state. These findings suggest that a pair of ES-responsive AstA-ergic neurons is required for inducing a stress-induced internal state.

### AstA-mediated action via its galanin receptor homolog AstA-R1 is required for stress-induced claustrophobia-like behavior

Two receptors, AstA-R1 and AstA-R2, have been identified to respond to AstA, which are preferentially expressed in the brain and gut, respectively ^36^. Knockdown of *AstA-R1* using the pan-neuronal GAL4, *Elav-GS* ^37^ that is activated by feeding flies RU486, impaired the development of claustrophobia-like behavior (Fig. 4a, Supplementary Fig. 4a). Furthermore, *AstA-R1*-knockdown after shock exposure suppressed the shock-induced claustrophobia-like behavior (Fig. 4b, Supplementary Fig. 4b). Therefore, AstA-R1 is required for induction and maintenance of the stress-induced claustrophobia-like behavior.

**Fig. 4.**
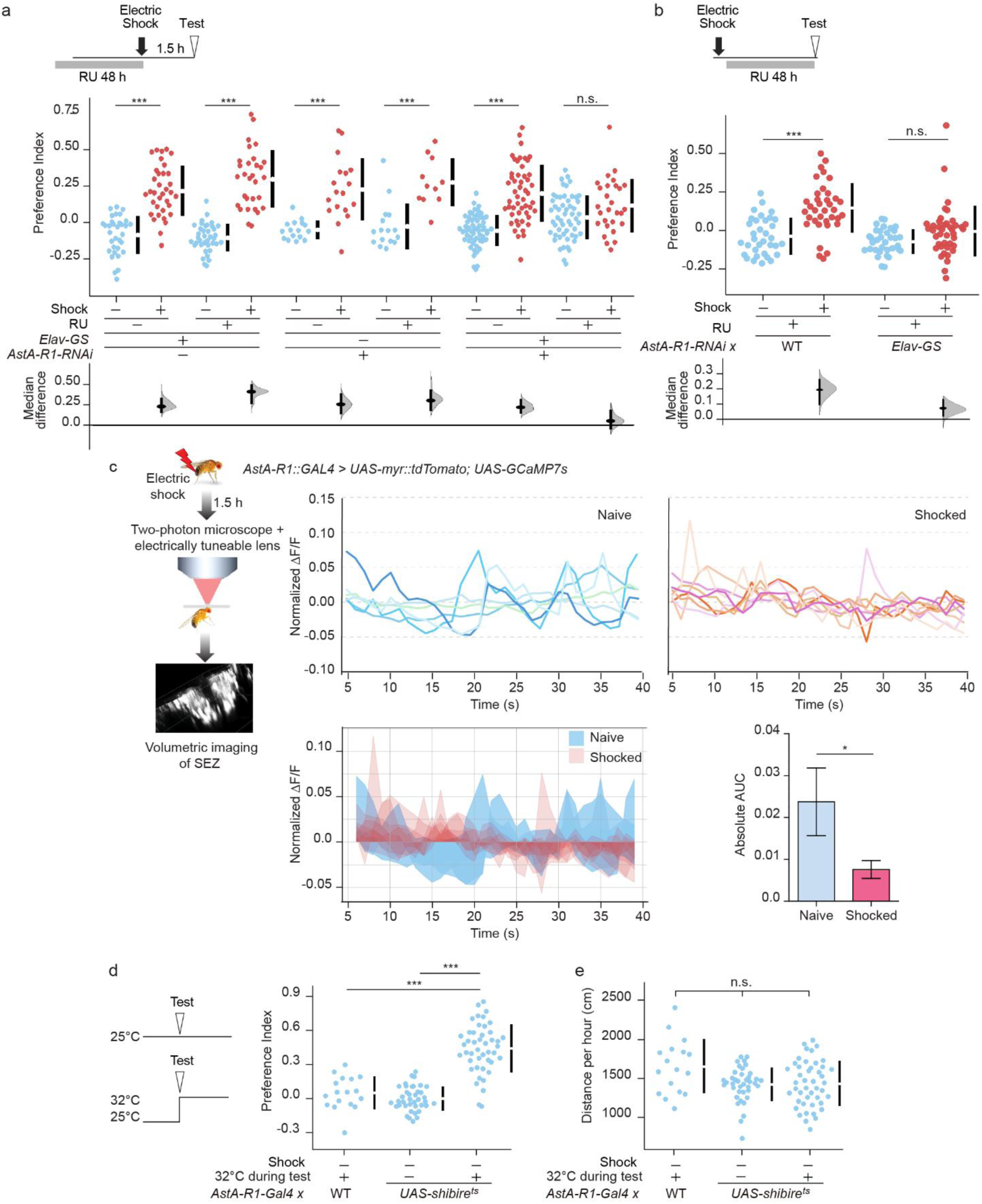
Inactivation of AstA-R1-expressing neurons is causally related to claustrophobia-like behavior. **a**,**b,** Knockdown of *AstA-R1* before or after shock exposure suppressed the claustrophobia-like behavior. The indicated flies were fed RU for 2 days before (a) or after electric shocks at 60 V for 5 min (b). The flies were rested for 1.5 h (a) or 2 d (b) and tested in the two-arm arena. Kruskal-Wallis test, a, *P* < 0.0001, *n* = 28-57; b, *P* < 0.0001, *n* = 34-41. **c,** Neural activity of AstA-R1-expressing neurons are reduced by electric shock. The indicated flies were subjected to electric shocks at 60 V for 5 min, rested for 1.5 h, and volumetric imaging was performed in the SEZ. The GCaMP7s signal traces were normalized to those of tdTomato and divided by the average of the total imaging duration (upper). The area under the curve (AUC) from time 5-40 sec was presented (lower left). Average absolute value from each fly was summarized in a graph (lower right). Two-sided Mann-Whitney U-test; *n* = 6, 8. **d,e,** Inactivation of AstA-R1 neurons induces claustrophobia-like behavior without shock exposure. The indicated flies were tested in the two-arm arena at 25°C or 32°C. Preference index (d) and locomotor activity indicated by total distance (cm) moved in the entire arena in 1 h (e) were shown. Kruskal-Wallis test, d, *P* < 0.0001, e, *P =* 0.073; *n* = 17-45. Data are represented as mean ± s.e.m., with median differences based on bootstrap estimation. n.s., not significant *P* > 0.05; *, *P* < 0.05; **, *P* < 0.01; ***, *P* < 0.001.

AstA-R1 is a receptor coupled to the G_i/o_ class of G proteins ^36^. Hence, we hypothesized that the shock-induced release of AstA suppresses the activity of AstA-R1-expressing neurons. We therefore conducted calcium imaging of the activity of AstA-R1-expressing neurons within the SEZ by expressing GCaMP7s and tdTomato using *AstA-R1-GAL4* in the naïve and shocked flies at 1.5 h after the shock. The direct comparison between naïve and shocked flies excludes the possibility that neural activity of AstA-R1-expressing neurons decays overtime during imaging. We captured a volumetric time series with voxel dimensions of approximately 0.5 × 0.5 × 3.6 µm, resulting in a total volume of 256 µm x 256 µm x ∼105 µm at a volume acquisition rate of 1 Hz for 40 sec, generating z-stuck series over time (Fig. 4c). The normalized GCaMP7s signal relative to tdTomato over the individual average demonstrated greater signal fluctuations in naïve flies more than in shocked flies (Fig. 4c). The absolute area under the curve (AUC) to quantify the integrated change of the normalized GCaMP7s signal revealed a significant reduction in the GCaMP7s signals after shock exposure (Fig. 4c). The reduction in activity of AstA-R1-expressing neurons may be causal to claustrophobia-like behavior. In fact, inactivation of AstA-R1 neurons during test by expressing *shibire^ts^* was sufficient to induce claustrophobia-like behavior without shock exposure (Fig. 4d), without affecting locomotor activity (Fig. 4e). Overall, these findings suggest that AstA-mediated inactivation of AstA-R1 expressing neurons is crucial for the development of the stress-induced internal state reflected in claustrophobia-like behavior.

### The Toll-signalling pathway in the perineurial barrier is involved in stress-induced claustrophobia-like behavior

Although the neuronal mechanism involving AstA signaling could be acute, flies gradually exhibit claustrophobia-like behavior over the course of 1.5-2 h post shock exposure (Fig. 5a, Supplementary Fig. 5a), suggesting another pathway with slower kinetics regulating the development of claustrophobia-like behavior, such as gene expression. In support of this idea, feeding cycloheximide (CHX), which blocks protein synthesis, impaired the development of claustrophobia-like behavior (Fig. 5b). To investigate the gene expression correlated to the manifestation of claustrophobia-like behavior, RNA sequencing (RNA-seq) was performed using the heads of individual naïve and shocked flies (Fig. 5c). The naïve flies with lower or higher PI were grouped as A or B, respectively, while the shocked flies with higher, medium, or lower PI were grouped as C, D, or E, respectively. Partial least squares discriminant analysis (PLS-DA) analysis determined that the segregation in gene expression between the groups was related to components 2 and 4 (Fig. 5d). To further characterize the gene expression change, each group of the shocked flies was compared with the naïve group. The increased genes were mostly found by comparing the naïve groups with the shocked group D that showed the medium degree of the claustrophobia-like behavior, but not with other shocked groups (Supplementary Fig. 5b). Accordingly, the shocked group D showed the broadest increase in gene expression compared to the other shocked groups, C and E (Supplementary Fig. 5c), indicating that change in gene expression appears in a subpopulation showing mild claustrophobia-like behavior in the shocked group. There were 29 overlapping genes whose expression was increased in the shocked group D compared to the native group (AB), the shocked group C, and E (Fig. 5e, 5f, Supplementary Table1), and their functions are related to Toll signaling and metabolism (Fig. 5g).

**Fig. 5.**
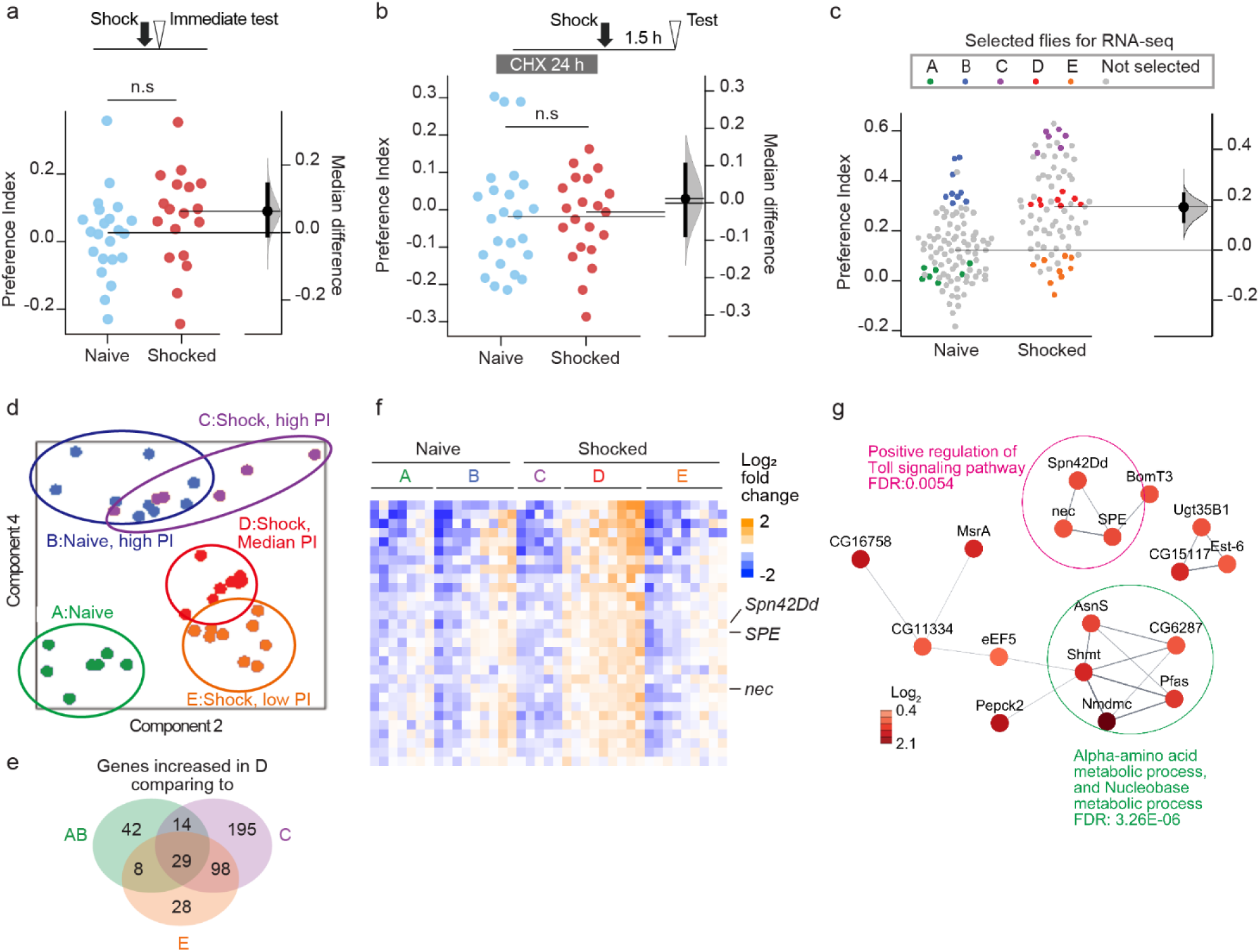
Shock exposure induces gene expression related to Toll signalling. **a**, The claustrophobia-like behavior is not observed immediately after shock exposure. The flies were subjected to electric shocks at 60 V for 5 min, and immediately tested in the two-arm arena for 1 h, Two-sided Mann-Whitney U-test; n = 18-22 **b,** Blocking protein synthesis impaired the development of claustrophobia-like behavior. The flies were fed 5 % sucrose with or without cycloheximide (CHX) for 24 h. The flies were subjected to electric shocks at 60 V for 5 min, rested for 1.5 h, and tested in the two-arm arena. Two-sided Mann-Whitney U-test; n = 21-24. **c,** Schematic of RNA-seq using individual fly heads. Fly heads were collected immediately after testing their behavior (approximately 2.5 h post-shock). **d,** PLS-DA analysis showing the segregation between the groups. **e,** Venn diagram indicating the overlaps between the increased genes. **f,** Heatmap of the log_2_ fold change in expression of 29 genes significantly increased in the shocked group D. **g,** Protein network within 29 genes identified from the STRING database. The log_2_ fold change of gene expression in the group D over that in the group A was shown. Data are represented as mean ± s.e.m., with median differences based on bootstrap estimation. n.s., not significant *P* > 0.05; *, *P* < 0.05; **, *P* < 0.01; ***, *P* < 0.001.

As activation of Toll signaling has been associated with behavioral change after social defeat in mice^12^, Toll signaling may be one of conserved mechanisms causally related to anxiety. The Toll signalling pathway (Fig. 6a) is essential for the *Drosophila* innate immune response to infections by fungi or Gram-positive bacteria through the activation of the transcription factor, Dorsal (*dl*) and Dorsal-related immunity factor (*Dif*), the *Drosophila* homolog of nuclear factor-kappaB (*NF-κB*)^38^. RNA-sequencing data demonstrated an increase in *Spätzle-Processing Enzyme* (*SPE*) expression (Fig. 5f, Supplementary Fig. 5d) which is required for processing pro-Spätzle (*pro-SPZ*) protein and producing the active Toll receptor ligand *SPZ* ^38^. In addition, *necrotic* (*nec*) and *Serpin 42Dd* (*Spn42Dd*), both involved in negative regulation of Toll signaling ^38^ also showed upregulation (Fig. 5f, Supplementary Fig. 5e, 5f), which could be related to the homeostatic control of Toll signaling. The change in gene expression after electric shock could be downstream of AstA-signaling. However, *AstA-R1*-knockdown did not reduce *SPE* expression (Supplementary Fig. 5g, 5h), suggesting that gene expression change is independent of AstA-signaling.

**Fig. 6.**
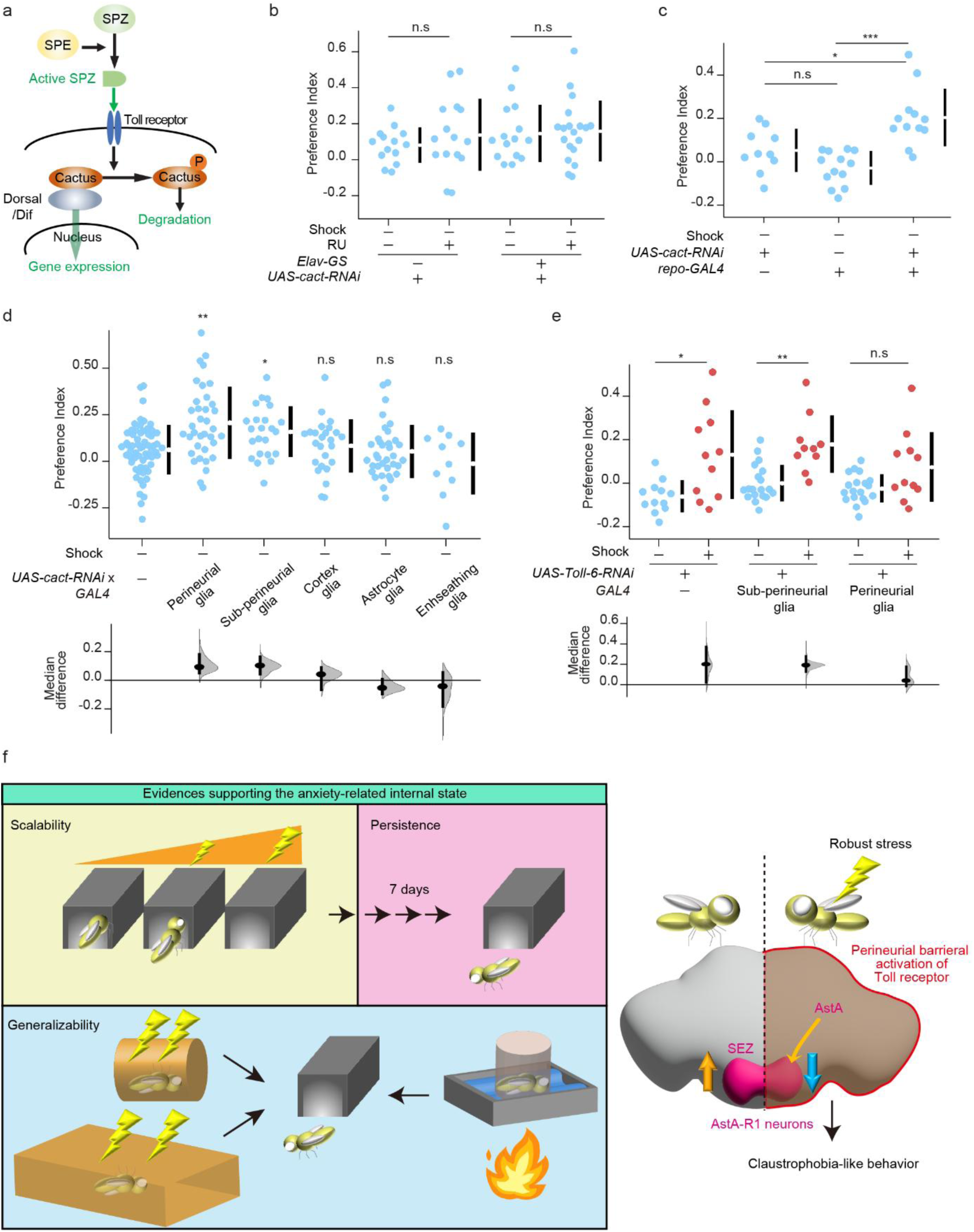
Toll-signalling pathway in perineurial glia is involved in developing claustrophobia-like behaviour. **a**, Schematic of Toll signaling pathway. **b-d,** Knockdown of *cactus*, not in neurons (b), but in glia (c), particularly in perineurial glia (d) induced claustrophobia-like behaviour without shock. The indicated flies were fed food containing RU for 2 days (b) or normal food (b,c,d). The naïve flies were tested in the two-arm arena. Kruskal-Wallis test, a, *P* = 0.5891, *n* = 14-19; b, *P =* 0.0002, *n* = 10-13; d, *P* < 0.0001, *n* = 10-61. **e**, Knockdown of *Toll-6* in perineurial glia, but not sub-perineurial suppressed the claustrophobia-like behaviour. The indicated flies were fed food containing RU for 2 days or normal food. The flies were subjected to electric shocks at 60 V for 5 min, rested for 1.5 h, and tested in the two-arm arena. Kruskal-Wallis test, *P* < 0.0001; *n* = 10-19. **f,** Model. Robust stress induces activations of AstA-releasing neurons and Toll signaling in perineurial glia, which act as variables generating the internal states causing claustrophobia-like behavior. Data are represented as mean ± s.e.m., with median differences based on bootstrap estimation. n.s., not significant *P* > 0.05; *, *P* < 0.05; **, *P* < 0.01; ***, *P* < 0.001.

Toll signaling can be genetically activated by knockdown of *cactus* (*cact*), which encodes a protein inhibiting Dorsal and Dif ^39^, the *Drosophila* homolog of inhibitory κB (IκB). We therefore tested whether knockdown of *cact* sufficiently induces claustrophobia-like behavior without applying electric shocks. While *cact*-knockdown in neurons by *Elav-GS* did not (Fig. 6b), that in glia by *repo-GAL4* induced the development of claustrophobia-like behavior (Fig. 6c). The *Drosophila* nervous system contains five subtypes of glial cells ^40^. Without subjecting flies to electric shock, claustrophobia-like behavior was induced significantly by knockdown of *cact* in perineurial glia and slightly by that in subperineurial glia (Fig. 6d, Supplementary Fig. 6a). Perineurial glia and sub-perineurial glia form the blood-brain barrier in *Drosophila* ^40^, where the inner layer of sub-perineurial glia creates the physical and chemical barrier, while the outer perineurial glia directly interfaces with the circulating haemolymph. *Drosophila* has nine Toll-receptors (Toll-1 to Toll-9), with Toll-6 and Toll-7 highly expressed in the central nervous system ^41^. Knockdown of *Toll-6* in glia (Supplementary Fig. 6b), but not neurons (Supplementary Fig. 6c) significantly reduced the claustrophobia-like behavior. Knockdown of *Toll-6* in perineurial glia suppressed the claustrophobia-like behavior, but that in sub-perineurial glia did not (Fig. 6e). Knockdown of *Toll-7* in neurons and perineurial glia did not affect the claustrophobia-like behavior (Supplementary Fig. 6d, 6e). Taken together, the Toll-6 receptor-mediated response in perineurial glia would be a slower component required for the development of claustrophobia-like behavior.

## Discussion

The active choices of animals during exploration can indicate internal states which could be related to anxiety. Research on fruit flies has primarily focused on aversive memory formation and its influence on future choices based on memory recall, with limited exploration of how stress exposure affects the broader behavioral repertoire. In this study, we utilized fruit flies to examine the after-effects of robust stress by introducing a behavioral assay assessing their active choice during exploration. This provided a unique opportunity to study the evolutionary origins of the generalized internal state potentially leading to anxiety.

Claustrophobia-like behavior could be one of the manifestations of the anxiety state, because it represents the proposed properties such as scalability, generalizability, and persistence, inferring the internal states ^26, 42^ (Fig. 6f). Scalability was supported by the elevated claustrophobia-like behavior according to intensities and durations of electric shock (Fig. 2a, 2b). Generalizability was indicated by the fact that claustrophobia-like behavior was induced by electric shock given in larger field (Fig. 2g) or by heat, a different modality of the valence (Fig. 2c). Persistence was demonstrated by the sustainability of individual degrees of claustrophobia-like behavior (Fig. 2d, 2e). Evoking fear by exposure to specific spaces is one of the hallmarks in phobias, as agoraphobia is related to an open space and claustrophobia is to a confined space. Ethologically, these features are rational to avoid inescapable situations from predators, which would involve a visual perception of the given environment. As flies in dark conditions did not display claustrophobia-like behavior (Supplementary Fig. 1), the visual perception of “confined” space may be targeted by the internal state related to anxiety. However, we determined that the claustrophobia-related AstA/AstA-R1 neurons reside in the SEZ where descending neurons important for motor outputs are localized, suggesting that AstA-signaling would modify the neural circuitry related to post-visual processing, such as decision-making in exploration or risk assessment, rather than primary visual processing in optic lobes as exemplified in modification of threat perception via dopamine ^43^. Importantly, manipulating AstA signaling did not affect reduced locomotion caused by shock exposure, suggesting that specific nodes related to active choices during exploration are biased by AstA-dependent pathway to accommodate the internal state, enhance risk assessment, and adopt the behavior to the given environment. Moreover, given that reduced locomotion or immobility is a hallmark of depression and manipulating AstA pathway did not affect locomotor activity, AstA signaling would orchestrate specific components related to anxiety, but not depression. AstA is also involved in feeding behavior^44^, appetitive learning^45^, water-seeking behavior^46^, and sleep ^47^, although its target neurons would be different. AstA may generally have modulatory functions in diverse types of behavior, depending on the target neurons.

We found that persistent inactive state of AstA-R1 expressing neurons is the neural correlates of claustrophobia-like behavior, mediated by the AstA-releasing neuron. AstA-R1 has sequence homology to the galanin receptor, suggesting that AstA/AstA-R1 pathway may be functionally conserved in the galanin/receptor pathway. Indeed, galanin in the locus coeruleus is required for long-lasting stress-induced anxiety ^7^. Activation of galanin-releasing LC neurons sufficiently induced the anxiety state ^6^. However, the actions and effects of galanin are complex ^48^ due to the receptor diversity. Galanin receptor 1 (GalR1) and 3 (GalR3) are Gi/Go-coupled receptors, whereas galanin receptor 2 (GalR2) is a Gq-coupled receptor ^49^. While GalR3-knockout mice show an anxiety-like phenotype ^50^, GalR2-knockout mice display reduced anxiety ^51^. The phenotype may vary depending on tests and methods, as galanin-releasing LC neurons target distinct brain regions, including the lateral septum and amygdala ^6^. More detailed molecular actions of AstA- and galanin-induced internal state would be of great interest. Importantly, claustrophobia-like behavior was suppressed by knocking down AstA-R1 after shock exposure, suggesting that the AstA-R1 functions after stress exposure are also significant to sustain the internal state. Further investigation of the AstA-R1 functions would provide more details of how the generalized internal states persist over time and may provide entry points to develop anxiolytic drugs.

Our data demonstrated the role of the perineurial barrier in the development of claustrophobia-like behavior. The perineurial barrier is a part of the blood-brain barrier, controlling the imports of sugar ^52^ and other substances ^53, 54^. Toll signaling activation in the perineurial barrier may lead to the change in transporter expression, which may eventually alter the internal environment of the brain. The Toll-6 downstream genes involved in this step would not be related to the known immune response genes, as neuronal Toll signaling required for larval locomotion did not involve canonical downstream genes in immune response, such as *Drosomycin* ^41^. The ligand regulating Toll-6 could be NT1 and NT2^41^, which may be circulating in the haemolymph. Importantly, NT1 and NT2 undergo proteolytic digestion via SPE, becoming the active ligand of Toll-6. Upregulation of *SPE* expression by shock exposure would have a causal role to induce Toll-6 activation. Paradoxically, we also found increased expression of negative regulators of Toll signaling, *nec* and *Spn42Dd*, suggesting the homeostatic control of Toll signaling after stress exposure which may allow local activation. The increase in *SPE* expression was detected only in the group displaying a mild degree of claustrophobia-like behavior. Thus, the immune response activation would be one of the biological variables determining the generalized internal state after stress. In mice, Toll-like receptors, TLR2 and TLR4 in microglia are involved in persistent social avoidance after repeated social defeat through activation of inflammation-related cytokines^12^. Although the downstream events controlled by Toll signaling may differ among cell types or animal species, the evolutional conservation and divergence of the Toll signaling pathway may suggest the traits regulating the generalized internal state.

Our findings connect stress-induced behaviors to neuronal and molecular representations of the internal state, taking advantage of *Drosophila* as a genetically tractable model organism to investigate anxiety. The relationship between the mechanisms involving AstA/AstA-R1 in neurons and Toll signaling in the perineurial barrier is yet to be determined. Both may be independently required for the development of the internal state or may be activated in parallel in different individuals, acting as variables determining the internal states (Fig. 6f). Further studies are needed to understand the complex interactions of mechanisms contributing to the stress-induced state and associated behavioral changes.

## Acknowledgements

We thank all lab members for their helpful discussions. We would like to thank Yi Rao, Ronald L Davis, Bloomington Drosophila Stock Center (NIH P40OD018537), FlyLight Project Team at Janelia Research Campus and Vienna Drosophila RNAi Center (VDRC, https://vdrc.at) for the materials. We also thank HKUST Biosciences Central Research Facility (BioCRF) team for their assistance. We thank Gewei Yan, Jianan Y. Qu, Xinyue Cui and Nilay Yapici for sharing information regarding fly preparation for calcium imaging, and Daniel Münch and Carlos Ribeiro for analyses using a two-photon microscope. AGA would like to thank Mohammad Farhan for helpful discussions. This study was supported by the JST PRESTO program. AGA work is supported by HKUST Postgraduate Studentship (PGS) and HKSAR Government Scholarship Fund - Belt and Road Scholarship.

## Author contributions statements

AGA and YH conceived the study and designed the experiments. AGA designed and conducted most of the behavioral experiments of the study. XH conducted most of the RNAi-based knockdown behavioral experiments following RNA-seq analysis. YG provided assistance with the behavioral setup. AGA performed the in-vivo calcium imaging experiments, with expert assistance from GT, JLS, KS, HT, SN, TM, and MS. YH conducted RNA sequencing library preparation. AGA assisted with RNA sequencing sample preparation and carried out the data analysis. KT helped with the RNA-seq data analysis. AGA and YH wrote the initial manuscript. YH and JLS revised the manuscript.

## Methods

### Fly strains

Our wild-type control line, w(CS10) ^55^ was used for most behavioral experiments or as a control unless otherwise stated. The *dumb2* mutant fly line ^17^ was obtained from M. Saitoe (Tokyo Metropolitan Institute of Medical Science, Japan). *Elav-GS* ^37^ and *MBsw* ^29^ was obtained from R. L. Davis (University of Florida, USA). All neuropeptide GAL4 of *AstA, Ms, Crz, Dsk, SIFa, PDF, sNPF, Trissin, CNMa, Trh, HDC-RC, Mip, NPF*, and *TK*, *AstA-R1-GAL4*, and their control line *w^1118^* were obtained from Yi Rao ^31^ (Peking University, China). *UAS-Kir2.1* ^28^ (6596), *UAS-IVS-myr::tdTomato* (32222)*, UAS-IVS-jGCaMP7s* ^33^ (79032), *Repo-GAL4* (7415)*, UAS-cactus-RNAi* (34775), and *GMR85G01-Gal4* (Perineurial glia-Gal4, 40436) lines were obtained from Bloomington Drosophila Stock Center (Indiana, USA)*. UAS-shibire^ts^ (pJFRC100-20XUAS-TTS-Shibire-ts1-p10),* split-GAL4 SS32423, spGAL4 control were from (Janelia Research Campus, U.S.A.). *UAS-AstA-R1-IR* (KK108648)*, UAS-Toll-6-RNAi* (GD928), *UAS-Toll-7-RNAi* (GD39176) were from Vienna *Drosophila* RNAi Center (Vienna, Austria). Subperineurial glia *GAL4* (NP2276, 112-853), cortex glia *GAL4* (NP2222, 112-830), ensheathing glia *GAL4* (NP6520, 105-240), and astrocyte-like glia *GAL4* (NP1243, 103-953) were from Kyoto DGGR ^56^.

### Fly culture conditions

Flies were raised under a 12-h light:dark cycle, at a temperature of 25 °C and humidity of 60%. Video-recordings for behavioral tests were conducted at the time of the evening peak, when locomotor activity is high. RU486 (RU; Mifepristone, Sigma, St. Louis, MO, USA) was dissolved in ethanol and mixed with fly food, to a final concentration of 0.5 mM RU, and flies were fed RU-containing food for 2 days prior to each experiment. For cycloheximide (CHX)-feeding, Whatman 3MM filter paper was soaked with a 5% sucrose solution containing 35 mM CHX, and flies were reared on the paper for 24 h.

### Treating flies with aversive stimuli

Flies were treated with 1.5-s pulses of electric shocks at 60 V every 5 s in total 5 min unless indicated. Heat treatment was performed by loading flies to the prewarmed plastic vials in the incubator set at the indicated temperature for 10 min. Vibration stress was performed by loading flies in 1.5 ml plastic tubes placed on a vortex mixer for 2.5 h. Restraint stress was performed by placing flies between plastic vials and soft sponge plugs for 2.5 h, which are generally used for fly culture.

### Recording of fly behavior

The two-arm arena was 3D-printed using white PLA material. Behavioral recordings were conducted inside a custom-made acrylic or metal box (24-25°C; ambient humidity) equipped with internal acoustic insulation to minimize surrounding noise. A Basler camera (acA1300-60gmNIR GigE), paired with a Kowa lens (LM12HC-SW), was mounted above the arena. White light was placed inside the metal box, while infrared (IR) light was illuminated as a backlight to track fly locomotion. An IR long-pass filter (MidOpt LP815-35.5) was placed on the camera lens to block the visible light. Flies were acclimated in the two-arm arenas for approximately 5-10 min before recording for 1 h. The recordings were captured at 15-30 frames per second using Noldus EthoVision-XT15 or MediaRecorder4.0 image acquisition software. The tracking of flies and behavior analyses was performed by EthoVision XT15. In the estimation graphs ^57^, we used a bootstrap estimation of effect sizes, plotting the data against a median difference using 5,000 bootstrapped resamples, each black dot indicates a median difference, and the associated black ticks depict error bars representing 95% confidence intervals; the shaded area represents the bootstrapped sampling-error.

### Behavioral data analysis

Analyses were done in R (https://www.r-project.org/, version 4.3.2), using the dabestr ^57^, plyr ^58^ and ggplot2 ^59^ packages.

### Olfactory aversive memory assay

Aversive olfactory conditioning was performed as previously described ^60, 61^. Briefly, approximately 70 flies were placed in a training chamber, where they were exposed to odors and electrical shocks. During single-cycle training, either 3-octanol (OCT, 218405, Sigma, St. Louis, MO, USA) or 4-methylcyclohexanol (MCH, 153095, Sigma, St. Louis, MO, USA) dissolved in mineral oil (O122-4, Thermo Fisher Scientific, San Jose, CA, USA) was paired with electrical shocks (60 V, 1.5-s pulses every 5 s) for 1 min, while the remaining odor was not. For testing, flies were placed at a choice point between the two odors for 1.5 min. A performance index (PI) was calculated so that a 50:50 distribution (no memory) yielded a PI of zero, while a 0:100 distribution away from the shock-paired odor yielded a PI of 100. Individual performance indices were calculated as the average of two experiments, in which the shock-paired odor was alternated.

### Immunohistochemistry

Staining of pERK was carried out as previously described^62^. Flies were briefly washed with ice-cold ethanol, which was then replaced with ice-cold phosphate buffered saline (PBS). All buffers below were supplemented with PhosSTOP (Sigma, St. Louis, MO, USA), with the exception of methanol. The proboscis was removed in ice-cold PBS within 5 min, following which the fly heads were fixed in 4% paraformaldehyde in PBS for 30 min on ice, followed by a second fixation with 100% methanol for 1 h on ice. Methanol was replaced with PBS, and the fixed brains were dissected. The dissected brains were treated with Proteinase K (Takara, Shiga, Japan) at 1 ug/ml in PBS containing 0.3% Triton X-100 for 30 min at room temperature (20-25°C), followed by post-fixation with 4% paraformaldehyde in PBS for 30 min at room temperature. The brains were washed three times with PBS and blocked for 1 h in PBS containing 0.1% Triton X-100 and 4% Block Ace (Sumitomo Dainippon Pharma, Osaka, Japan). The samples were then incubated with primary antibodies in blocking solution for 1 d at 4°C. After washing three times with PBS containing 0.1% Triton X-100, the brains were incubated with secondary antibodies in blocking solution for 6 h at 4°C. After washing three times with PBS containing 0.1% Triton X-100, the brains were mounted in PermaFluor (Lab Vision Corp., Fremont, CA, USA), and images were captured using a confocal microscope, LSM980 (Zeiss Microsystems, Jena, Germany). The following primary antibodies were used: in Fig. 3c, mouse monoclonal anti-pERK (M8159, Sigma, 1:100) and rabbit anti-DsRed (632496, Takara Bio, 1:100); in Supplementary Fig. 3c, rabbit monoclonal anti-pERK (4376, Cell Signaling Technology, 1:100), mouse monoclonal anti-Diploptera punctata Ast7 (5F10, DSHB, 1:4, recognizing AstA in *Drosopihila*), and rat anti-tdTomato (EST203, Kerafast, 1:100). The following secondary antibodies were used: donkey anti-mouse IgG Alexa Fluor 488 (715-545-150, Jackson ImmunoResearch Labs), Goat anti-rabbit IgG Alexa Fluor 555 (A-21429, Invitrogen), Goat anti-Rabbit IgG Alexa Fluor 488 (A-11034, Invitrogen), Goat anti-Mouse IgG Alexa Fluor 647 (A-21236, Invitrogen) and Donkey anti-rat IgG Cy3 (712-165-153, Jackson, ImmunoResearch Labs). Secondary antibodies were diluted to 1:400 in blocking solution, together with DAPI at 2.5 ug/ml.

### Fly preparation for two-photon imaging

Six- to seven-day old male flies were briefly cold anesthetized (for <30 s) and then tethered to a piece of tape using a human hair placed across the neck ^63^. The proboscis was glued in a partially extended position using UV-curable glue (Bondic). Then the tape was attached to a metal piece with a hole of 7 mm-diameter to accommodate the fly body. A small hole was cut into the tape, precisely above the head, to allow the top of the head capsule to extend above the plane of the tape. A dot of UV-curable glue was applied to the eyes and the neck to restrict head movement. Then a drop of Adult Haemolymph-Like Saline (AHLS) (108 mM NaCl, 5 mM KCl, 2 mM CaCl2, 8.2 mM MgCl2, 4 mM NaHCO3, 1 mM NaH2PO4, 5 mM trehalose, 10 mM sucrose, 5 mM HEPES, pH7.5, osmolarity adjusted to 270 mOsm) was added to cover the head capsule which was then opened by carefully cutting and folding back the flap of cuticle covering the dorsal-anterior portion of the head, including the antennae. Trachea and fat covering the brain were removed and the esophagus was transected to allow for unoccluded visual access to the SEZ ^64^. In the case of delivering foot shock while simultaneously conducting two-photon imaging, the metal piece holding the fly was then positioned on a PLA custom-made 3D-printed platform designed to accommodate a glass slide coated with a transparent and conductive indium tin oxide (ITO) circuit, connected to the electric shock delivery system via two electrodes. The platform was designed to allow for ∼1mm distance between the fly abdomen and the glass slide, allowing for the fly legs to touch the electric grid and receive the electric shocks.

### Two-photon imaging

In-vivo calcium imaging was performed using a two-photon microscope (Nikon A1R MP+) equipped with a 25× water-immersion objective (Nikon CFI75 Apochromat 25XC W 1300, 1.1 numerical aperture). The excitation laser was calibrated to a wavelength of 920 nm. For imaging the AstA-GAL4 response to electric shock (Fig. 3d), recordings were taken at the same plane at 30 frames per second for 210 sec, with a resolution of 512×512 (0.33 μm/pixel). For SEZ volumetric imaging of AstAR1-GAL4, an electrically tuneable lens (Optotune EL-16-40-TC) mounted behind the objective was used to enable multi-plane imaging. Volumetric imaging involved capturing 30 Z-planes at a volume acquisition rate of 1 Hz over a duration of 40 sec. The voxel dimensions were 0.5 µm x 0.5 µm x 3.6 µm, resulting in a total volume of 256 µm x 256 µm x ∼105 µm. Imaging of the shocked flies took place approximately 90 min after shock exposure (the time by which claustrophobia-like behavior is manifested). A total of 6 naïve flies and 8 shocked flies were imaged.

### Image Processing

Image processing was performed using FIJI/ImageJ ^65^. In Fig. 3d, data acquisition took place at the same plane for 210 sec, structured as three alternating rounds: each round consisted of a 30-sec resting period followed by an intermittent shock episode lasting a total of 30 sec (Fig. 3e), concluding with another 30-sec rest. Fluorescence intensity was calculated by manually drawing a region of interest within the SEZ region. For F_0_ calculation, the average signal of 10 sec (300 frames) within the resting period was calculated for both GCaMP7s and tdTomato, then 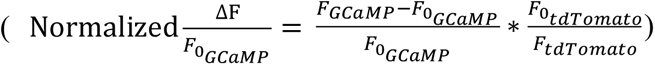 was calculated, and plotted using ImageJ. In the case of volumetric imaging, recording was done for 40 sec, excluding the first 5 sec from analysis due to photobleaching. Then fluorescence intensity was calculated by manually drawing a region of interest around the SEZ region in the Z-projection of the volume collected at each time point. Avg(*F*_*GCaMP*_) was calculated for the entire duration for GCaMP7s and tdTomato signal, then 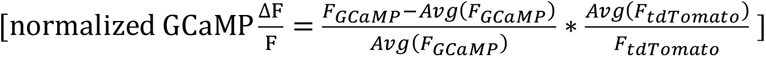 was calculated. The absolute Area Under Curve was the estimated by Simpson’s Rule using python v.3.10.16 (https://www.python.org/downloads/release/python-31016/), and python packages scipy^99^, numpy^100^ and pandas^101^.

### RNA sequencing

Individual heads were collected and homogenized using TRIzol reagent (Thermo Fisher Scientific, San Jose, CA, USA). RNA was extracted with chloroform, ethanol-precipitated, and dissolved in 6 µL diethylpyrocarbonate (DEPC) water. DNA was digested with DNase, and mRNA was further collected using Oligo d(T)25 Magnetic Beads (S1419, New England Biolabs inc., Ipswich, MA, USA), according to the manufacturer’s instructions. The obtained mRNA was then used to generate a library with the NEBNext Ultra II RNA Library Prep Kit for Illumina (New England Biolabs Inc., Ipswich, MA, USA), according to the manufacturer’s instructions. Paired-end reads were generated on a HiSeq X Ten (Illumina, San Diego, CA, USA).

### Bioinformatic analysis

The quality of the sequencing reads was initially evaluated using FastQC (version 0.11.4) (http://www.bioinformatics.babraham.ac.uk/projects/fastqc/), and proceeded to adaptor trimming using Trimmomatic ^66^, followed by mapping to the *Drosophila* reference genome, dm6, from UCSC using STAR ^67^. Reads with mapping quality below eight and non-primary mapped reads were eliminated. The filtered reads (5.4-12.3 million reads) were analyzed with HTSeq-count ^68^ to obtain the number of reads mapped on exons, which were further analyzed and normalized on R ^69^ using DESeq2 ^70^. To extract gene expression variations between groups hidden within noisy RNA-Seq data and visualize the resulting low-dimensional representation of samples, dimensionality reduction was performed using PLS-DA ^71^ implemented in the guidedPLS package (https://cran.r-project.org/web/packages/guidedPLS). The relative rpkm value was visualized and color-coded using Java TreeView ^72^.

### Quantification of transcripts (RT-qPCR)

Total RNA was extracted from 20 heads using TRIzol reagent (Thermo Fisher Scientific, San Jose, CA, USA), and 125 μg of RNA was used to synthesize cDNA using ReverTra Ace qPCR RT Master Mix with gDNA Remover (TOYOBO, Osaka, Japan). The cDNAs were then analyzed via quantitative real-time PCR (BioRad Laboratories, Hercules, CA, USA). Transcripts of *gaph2* were used for normalization.

### Statistical analysis

No statistical calculations were used to predetermine sample sizes. Our sample size was similar to those generally used in this field of research. Flies from each cross were randomly assigned to the treatment groups where possible. For behavioral experiments, all samples were numbered, and the investigators were blinded. For other experiments, blinding was not performed. Statistical analyses were performed using Prism version 9. For behavioral data analysis, the Mann-Whitney U test was used for comparisons between two groups, and the Kruskal-Wallis test, followed by Dunn’s multiple comparison test, was used for comparisons among multiple groups. For qPCR data analysis, one-way ANOVA followed by Tukey’s post hoc test was used for comparisons among multiple groups. *P* values < 0.05 were regarded as statistically significant. All data are presented as mean ± s.e.m. Each experiment was successfully reproduced at least twice and performed on multiple days.

## Data availability

The GEO accession number for the RNA-seq data reported in this paper is GSE294309.

---

**Supplementary Fig. 1.**
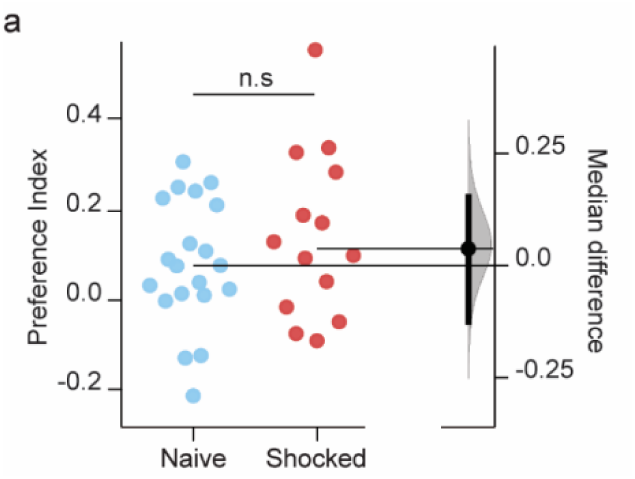
Claustrophobia-like behavior does not appear in the dark. Flies were shocked at 60 V, rested for 1.5 h, and tested in the two-arm arena in the dark. Two-sided Mann-Whitney U-test; n = 14, 20. Data are represented as mean ± s.e.m., with median differences based on bootstrap estimation. n.s., not significant *P* > 0.05.

**Supplementary Fig. 2.**
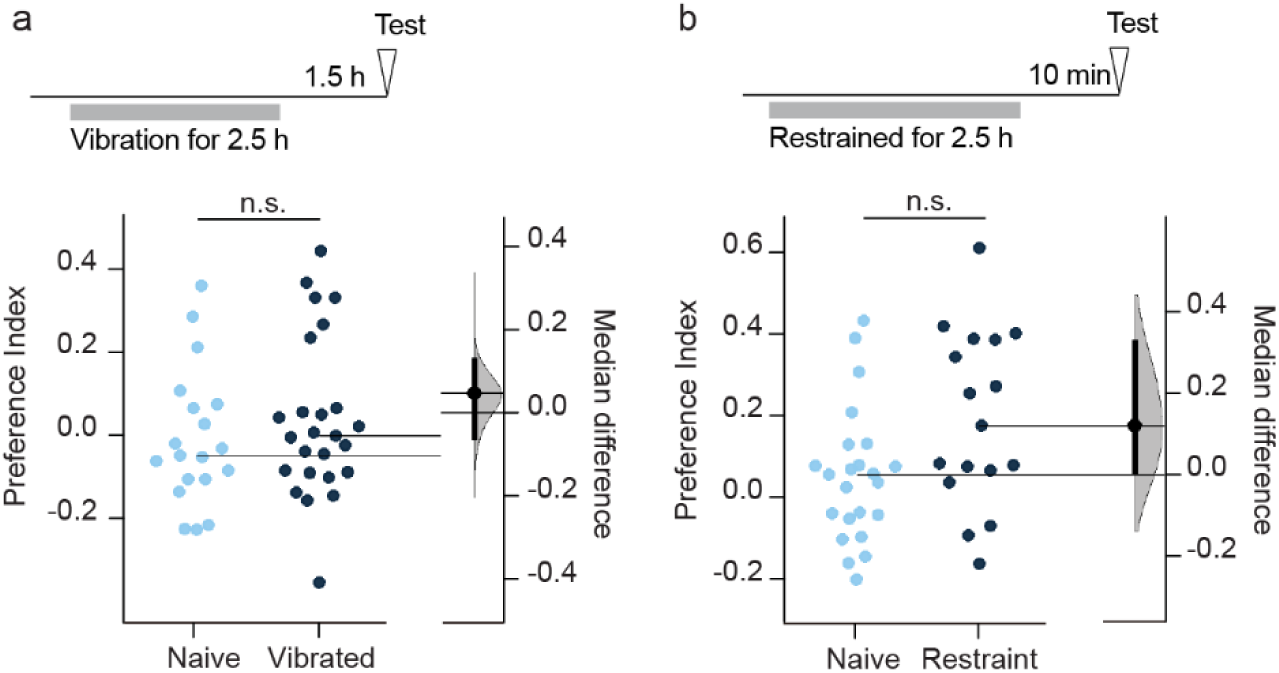
Vibration or restraint stress does not induce claustrophobia-like behavior. **a**, Vibration stress does not induce claustrophobia-like phenotype. Flies were exposed to vibrations for 2.5 h, rested for 1.5 h, and tested in the two-arm arena. Two-sided Mann-Whitney U-test; n = 17-23. **b,** Restraint stress does not induce claustrophobia-like behavior. Flies were restrained for 2.5 h, rested for 10 min, and tested in the two-arm arena. Two-sided Mann-Whitney U-test; n = 19-25. Data are represented as mean ± s.e.m., with median differences based on bootstrap estimation. n.s., not significant *P* > 0.05; *, *P* < 0.05; **, *P* < 0.01; ***, *P* < 0.001.

**Supplementary Fig. 3.**
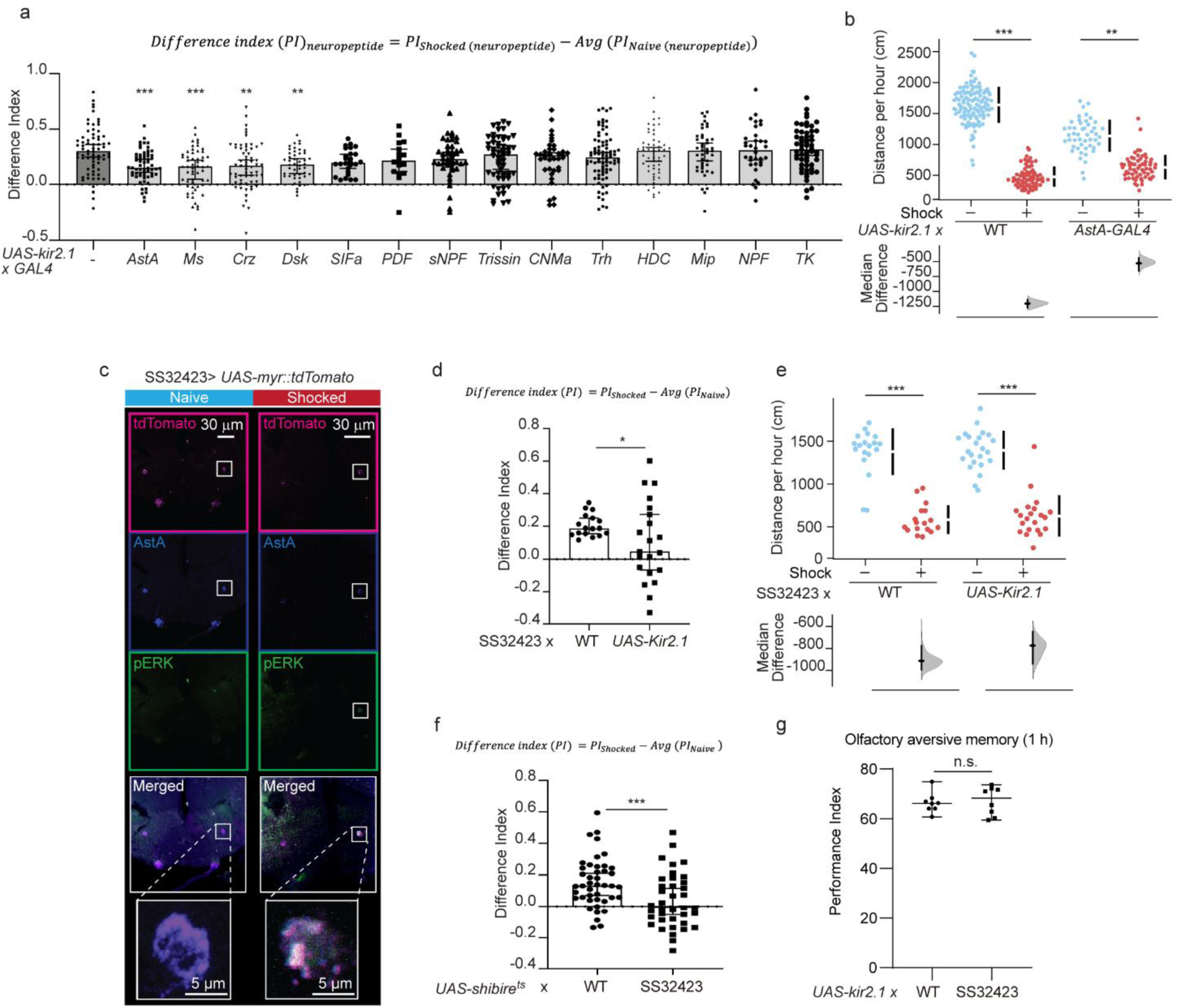
A pair of Allatostatin-A neuropeptidergic neurons is required for stress-induced claustrophobia-like behavior. **a,d,f**, The changes of the preference index after shock exposure. Differential indices were calculated by subtracting each preference index (PI) by average of PI of the corresponding naïve flies, which was presented in Fig. 3a, 3e, and 3f, respectively. a, Kruskal-Wallis test, *P* < 0.0001; *n* = 18-82. d, Two-sided Mann-Whitney U-test; n = 17-23. f, Two-sided Mann-Whitney U-test; n = 36-44. **b,** Silencing *AstA-GAL4*-labelled neurons partially alleviated the reduced locomotor activity in the shocked flies. The indicated flies were shocked at 60 V, rested for 1.5 h, and tested in the two-arm arena. Kruskal-Wallis test, *P* < 0.0001; *n* = 48-112. **c,** SS32423 split-GAL4 labels a pair of ES-responsive AstA-releasing neurons. The indicated flies were shocked at 60 V for 5 min. The brains were fixed 10 min after the shock and stained by anti-pERK, anti-tdTomato, and anti-AstA antibodies. The images are representative of three experimental replicates. **e,** Silencing the neurons labelled by SS32423 did not affect the reduction in the locomotor activity after shock exposure. The indicated flies were shocked at 60 V, rested for 1.5 h, and tested in the two-arm arena. Kruskal-Wallis test, *P* < 0.0001; *n* = 17-23. **g,** Silencing the neurons labelled by SS32423 did not affect olfactory aversive memory. The indicated flies were subjected to the olfactory aversive training and tested at 1 h after the training. Two-sided Mann-Whitney U-test; n = 8, 8. Data are represented as mean ± s.e.m., with median differences based on bootstrap estimation. n.s., not significant *P* > 0.05; *, *P* < 0.05; **, *P* < 0.01; ***, *P* < 0.001.

**Supplementary Fig. 4.**
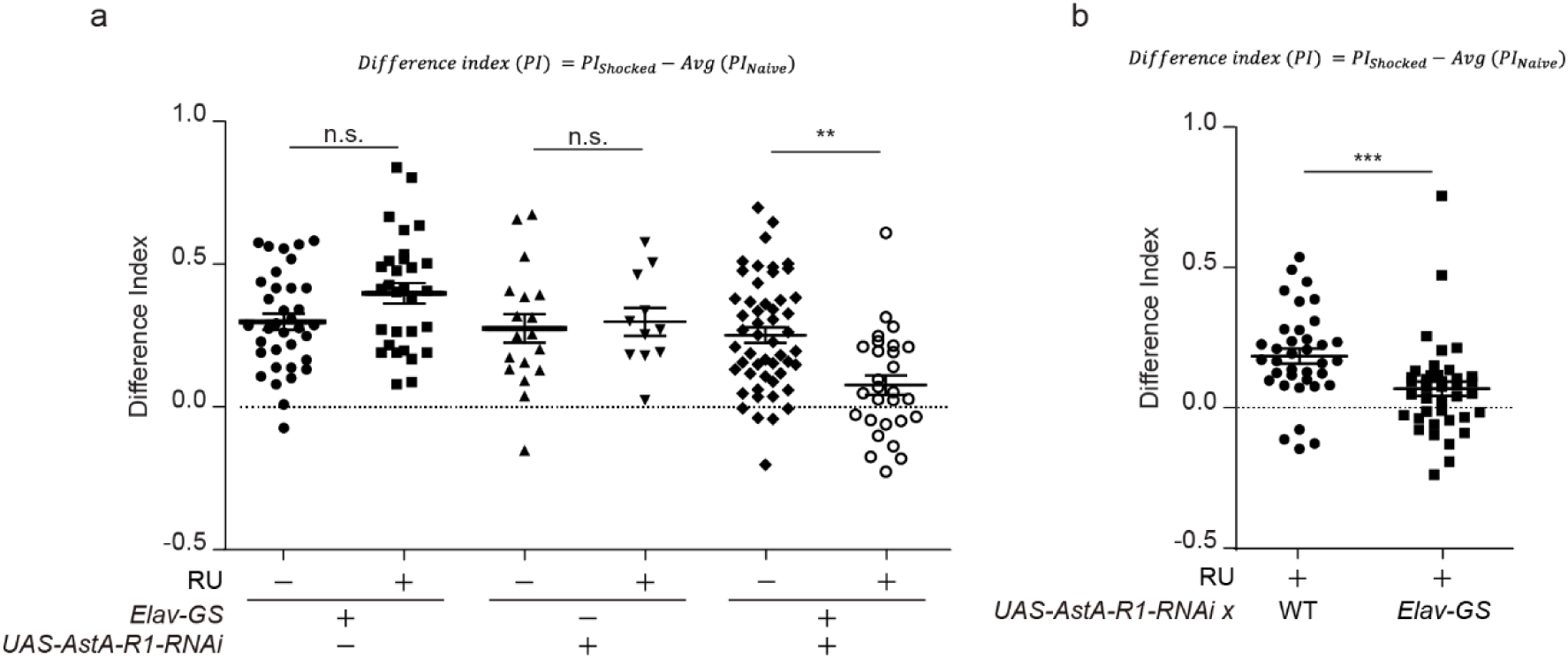
A pair of Allatostatin-A neuropeptidergic neurons is required for stress-induced claustrophobia-like behavior. **a,b,** The changes of the preference index after shock exposure in AstA-R1 knockdown flies. Differential indices were calculated by subtracting each preference index (PI) by average of PI of the corresponding naïve flies, which was presented in Fig. 4a and 4b, respectively. Two-sided Mann-Whitney U-test; a, *n* = 28-57; b, *n* = 34-41.

**Supplementary Fig. 5.**
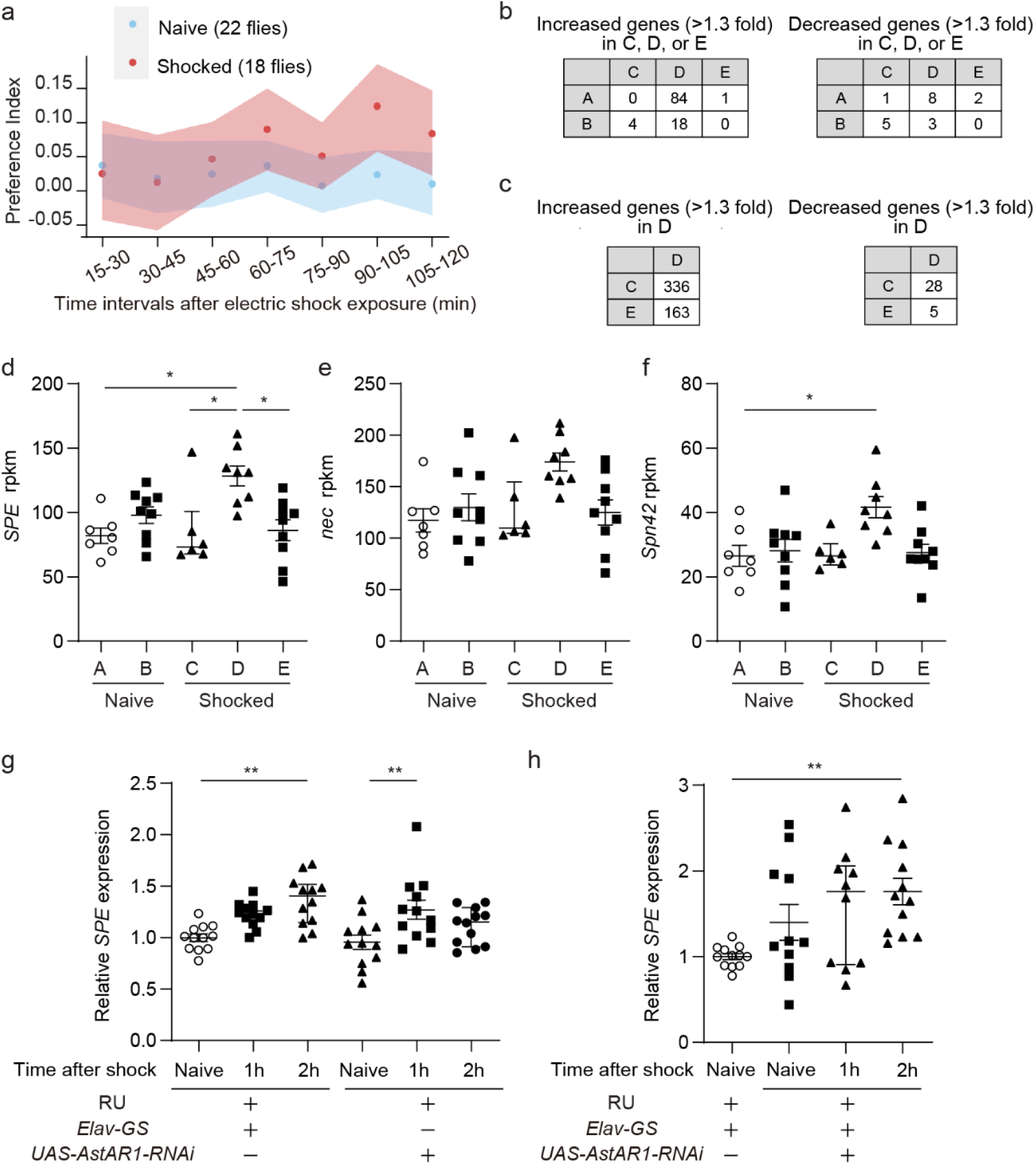
Shock exposure induces changes in gene expression. **a,** Shock exposure gradually induces claustrophobia-like behavior over the course of 1.5-2 h. Flies were subjected to electric shocks at 60 V for 5 min, and immediately analyzed in the two-arm arena. The dot and blanket indicated the median and the 85 % quantile, respectively. **b,c,** Differentially expressed genes among each group. The numbers of genes identified as increased or decreased genes showing *q* < 0.05 with fold change over 1.3 were presented. **d-f,** Relative gene expression of *SPE*, *nec*, and *Spn42Dd* in each group. Kruskal-Wallis test, d, *P* = 0.0065, *n* = 7-9; e, *P* = 0.0369, *n* = 7-9; f, *P* = 0.0199, *n* = 7-9. **g,h,** Shock exposure induces *SPE* gene expression independently of AstA-R1. One-way ANOVA, g, *P* < 0.0001, *n* = 12 for all; h, *P* = 0.012, *n* = 10-12.

**Supplementary Fig. 6.**
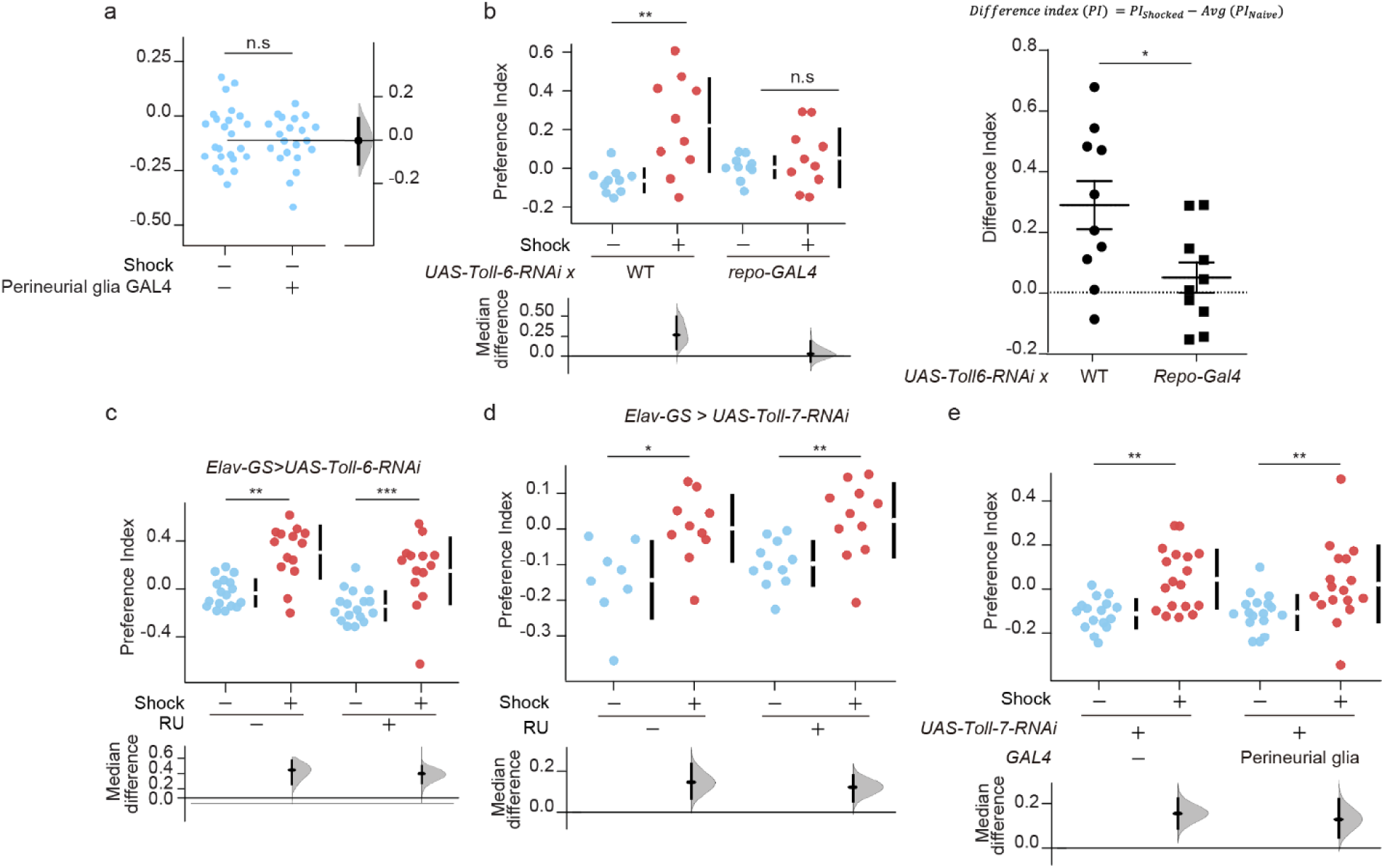
The involvement of Toll signaling in claustrophobia-like behavior. **a,** The genetic control flies carrying perineurial glial GAL4 does not affect the development of claustrophobia-like behavior. The indicated flies were subjected to electric shocks at 60 V for 5 min, rested for 1.5 h, and tested in the two-arm arena. Two-sided Mann-Whitney U-test; n = 21-24. **b,c,** The claustrophobia-like behavior is suppressed by knockdown of *Toll-6* in glia (b), but not in neurons (c). The indicated flies were subjected to electric shocks at 60 V for 5 min, rested for 1.5 h, and tested in the two-arm arena. Kruskal-Wallis test, b, *P* = 0.0086, *n* = 10 for all; c, *P* < 0.0001, *n* = 14-17. **d,e,** The claustrophobia-like behavior is unaffected by knockdown of *Toll-7* in neurons (d) or glia (e). The indicated flies were subjected to electric shocks at 60 V for 5 min, rested for 1.5 h, and tested in the two-arm arena. Kruskal-Wallis test, d, *P* = 0.0016, *n* = 8; *P* = 0.0003, *n* = 16-18. Data are represented as mean ± s.e.m., with median differences based on bootstrap estimation. n.s., not significant *P* > 0.05; *, *P* < 0.05; **, *P* < 0.01; ***, *P* < 0.001.

## References

1. Bienvenu TCM, Dejean C, Jercog D, Aouizerate B, Lemoine M, Herry C. The advent of fear conditioning as an animal model of post-traumatic stress disorder: Learning from the past to shape the future of PTSD research. Neuron 109, 2380–2397 (2021).

2. Dunsmoor JE, Cisler JM, Fonzo GA, Creech SK, Nemeroff CB. Laboratory models of post-traumatic stress disorder: The elusive bridge to translation. Neuron 110, 1754–1776 (2022).

3. King NJ, Eleonora G, Ollendick T. Etiology of childhood phobias: current status of Rachman’s three pathways theory. Behav Res Ther 36, 297–309 (1998).

4. Calhoon GG, Tye KM. Resolving the neural circuits of anxiety. Nature neuroscience 18, 1394–1404 (2015).

5. Botta P, et al. Regulating anxiety with extrasynaptic inhibition. Nature neuroscience 18, 1493–1500 (2015).

6. McCall JG, et al. CRH Engagement of the Locus Coeruleus Noradrenergic System Mediates Stress-Induced Anxiety. Neuron 87, 605–620 (2015).

7. Tillage RP, Foster SL, Lustberg D, Liles LC, McCann KE, Weinshenker D. Co-released norepinephrine and galanin act on different timescales to promote stress-induced anxiety-like behavior. Neuropsychopharmacology : official publication of the American College of Neuropsychopharmacology 46, 1535–1543 (2021).

8. Anthony TE, Dee N, Bernard A, Lerchner W, Heintz N, Anderson DJ. Control of stress-induced persistent anxiety by an extra-amygdala septohypothalamic circuit. Cell 156, 522–536 (2014).

9. Fuzesi T, Daviu N, Wamsteeker Cusulin JI, Bonin RP, Bains JS. Hypothalamic CRH neurons orchestrate complex behaviours after stress. Nature communications 7, 11937 (2016).

10. Niu M, et al. Claustrum mediates bidirectional and reversible control of stress-induced anxiety responses. Science advances 8, eabi6375 (2022).

11. Zhang GW, et al. Medial preoptic area antagonistically mediates stress-induced anxiety and parental behavior. Nature neuroscience 24, 516–528 (2021).

12. Nie X, et al. The Innate Immune Receptors TLR2/4 Mediate Repeated Social Defeat Stress-Induced Social Avoidance through Prefrontal Microglial Activation. Neuron 99, 464–479 e467 (2018).

13. Wohleb ES, Patterson JM, Sharma V, Quan N, Godbout JP, Sheridan JF. Knockdown of interleukin-1 receptor type-1 on endothelial cells attenuated stress-induced neuroinflammation and prevented anxiety-like behavior. The Journal of neuroscience : the official journal of the Society for Neuroscience 34, 2583–2591 (2014).

14. Zelikowsky M, et al. The Neuropeptide Tac2 Controls a Distributed Brain State Induced by Chronic Social Isolation Stress. Cell 173, 1265–1279 e1219 (2018).

15. Cognigni P, Felsenberg J, Waddell S. Do the right thing: neural network mechanisms of memory formation, expression and update in Drosophila. Current opinion in neurobiology 49, 51–58 (2017).

16. Guven-Ozkan T, Davis RL. Functional neuroanatomy of Drosophila olfactory memory formation. Learn Mem 21, 519–526 (2014).

17. Kim YC, Lee HG, Han KA. D1 dopamine receptor dDA1 is required in the mushroom body neurons for aversive and appetitive learning in Drosophila. The Journal of neuroscience : the official journal of the Society for Neuroscience 27, 7640–7647 (2007).

18. Ries AS, Hermanns T, Poeck B, Strauss R. Serotonin modulates a depression-like state in Drosophila responsive to lithium treatment. Nature communications 8, 15738 (2017).

19. Mohammad F, et al. Ancient Anxiety Pathways Influence Drosophila Defense Behaviors. Current biology : CB 26, 981–986 (2016).

20. Kim YK, et al. Repetitive aggressive encounters generate a long-lasting internal state in Drosophila melanogaster males. Proceedings of the National Academy of Sciences of the United States of America 115, 1099–1104 (2018).

21. Shohat-Ophir G, Kaun KR, Azanchi R, Mohammed H, Heberlein U. Sexual deprivation increases ethanol intake in Drosophila. Science 335, 1351–1355 (2012).

22. Felix RC, et al. Unravelling the Evolution of the Allatostatin-Type A, KISS and Galanin Peptide-Receptor Gene Families in Bilaterians: Insights from Anopheles Mosquitoes. PloS one 10, e0130347 (2015).

23. Karlsson R, Holmes A. Galanin as a modulator of anxiety and depression and a therapeutic target for affective dise. Amino Acids 31, 231–239 (2006).

24. Lewin BD. Claustrophobia. The Psychoanalytic Quarterly 4, 227–233 (1935).

25. Seong KH, et al. Paternal restraint stress affects offspring metabolism via ATF-2 dependent mechanisms in Drosophila melanogaster germ cells. Communications biology 3, 208 (2020).

26. Flavell SW, Gogolla N, Lovett-Barron M, Zelikowsky M. The emergence and influence of internal states. Neuron 110, 2545–2570 (2022).

27. Tully T, Preat T, Boynton SC, Del Vecchio M. Genetic dissection of consolidated memory in Drosophila. Cell 79, 35–47 (1994).

28. Baines RA, Uhler JP, Thompson A, Sweeney ST, Bate M. Altered electrical properties in Drosophila neurons developing without synaptic transmission. The Journal of neuroscience : the official journal of the Society for Neuroscience 21, 1523–1531 (2001).

29. Mao Z, Roman G, Zong L, Davis RL. Pharmacogenetic rescue in time and space of the rutabaga memory impairment by using Gene-Switch. Proceedings of the National Academy of Sciences of the United States of America 101, 198–203 (2004).

30. Nassel DR, Winther AM. Drosophila neuropeptides in regulation of physiology and behavior. Progress in neurobiology 92, 42–104 (2010).

31. Deng B, et al. Chemoconnectomics: Mapping Chemical Transmission in Drosophila. Neuron 101, 876–893 e874 (2019).

32. Li Q, et al. Importin-7 mediates memory consolidation through regulation of nuclear translocation of training-activated MAPK in Drosophila. Proceedings of the National Academy of Sciences of the United States of America 113, 3072–3077 (2016).

33. Dana H, et al. High-performance calcium sensors for imaging activity in neuronal populations and microcompartments. Nature methods 16, 649–657 (2019).

34. Meissner GW, et al. A searchable image resource of Drosophila GAL4 driver expression patterns with single neuron resolution. eLife 12, (2023).

35. Kitamoto T. Conditional modification of behavior in Drosophila by targeted expression of a temperature-sensitive shibire allele in defined neurons. Journal of neurobiology 47, 81–92 (2001).

36. Birgul N, Weise C, Kreienkamp HJ, Richter D. Reverse physiology in drosophila: identification of a novel allatostatin-like neuropeptide and its cognate receptor structurally related to the mammalian somatostatin/galanin/opioid receptor family. The EMBO journal 18, 5892–5900 (1999).

37. Osterwalder T, Yoon KS, White BH, Keshishian H. A conditional tissue-specific transgene expression system using inducible GAL4. Proceedings of the National Academy of Sciences of the United States of America 98, 12596–12601 (2001).

38. Valanne S, Wang JH, Ramet M. The Drosophila Toll signaling pathway. J Immunol 186, 649–656 (2011).

39. Drier EA, Huang Lh Fau - Steward R, Steward R. Nuclear import of the Drosophila Rel protein Dorsal is regulated by phosphorylation. Genes Dev 13, 556–568 (1999).

40. Kremer MC, Jung C, Batelli S, Rubin GM, Gaul U. The glia of the adult Drosophila nervous system. Glia 65, 606–638 (2017).

41. McIlroy G, et al. Toll-6 and Toll-7 function as neurotrophin receptors in the Drosophila melanogaster CNS. Nature neuroscience 16, 1248–1256 (2013).

42. Anderson DJ, Adolphs R. A framework for studying emotions across species. Cell 157, 187–200 (2014).

43. Cazale-Debat L, et al. Mating proximity blinds threat perception. Nature 634, 635–643 (2024).

44. Hergarden AC, Tayler TD, Anderson DJ. Allatostatin-A neurons inhibit feeding behavior in adult Drosophila. Proceedings of the National Academy of Sciences of the United States of America 109, 3967–3972 (2012).

45. Yamagata N, Hiroi M, Kondo S, Abe A, Tanimoto H. Suppression of Dopamine Neurons Mediates Reward. PLoS biology 14, e1002586 (2016).

46. Landayan D, Wang BP, Zhou J, Wolf FW. Thirst interneurons that promote water seeking and limit feeding behavior in. eLife 10, (2021).

47. Chen JT, et al. Allatostatin A Signalling in Regulates Feeding and Sleep and Is Modulated by PDF. Plos Genet 12, (2016).

48. Pérez de la Mora M, et al. Dysfunctional Heteroreceptor Complexes as Novel Targets for the Treatment of Major Depressive and Anxiety Disorders (2022).

49. Webling KE, Runesson J, Bartfai T, Langel U. Galanin receptors and ligands. Frontiers in endocrinology 3, 146 (2012).

50. Brunner SM, et al. GAL3 receptor KO mice exhibit an anxiety-like phenotype. Proceedings of the National Academy of Sciences of the United States of America 111, 7138–7143 (2014).

51. Bailey KR, Pavlova MN, Rohde AD, Hohmann JG, Crawley JN. Galanin receptor subtype 2 (GalR2) null mutant mice display an anxiogenic-like phenotype specific to the elevated plus-maze. Pharmacology, biochemistry, and behavior 86, 8–20 (2007).

52. Volkenhoff A, Weiler A, Letzel M, Stehling M, Klämbt C, Schirmeier S. Glial Glycolysis Is Essential for Neuronal Survival in *Drosophila*. Cell Metabolism 22, 437–447 (2015).

53. Seabrooke S, O’Donnell MJ. Oatp58Dc contributes to blood-brain barrier function by excluding organic anions from the Drosophila brain. American journal of physiology Cell physiology 305, C558–567 (2013).

54. Parkhurst SJ, Adhikari P, Navarrete JS, Legendre A, Manansala M, Wolf FW. Perineurial Barrier Glia Physically Respond to Alcohol in an Akap200-Dependent Manner to Promote Tolerance. Cell reports 22, 1647–1656 (2018).

55. Dura JM, Preat T, Tully T. Identification of linotte, a new gene affecting learning and memory in Drosophila melanogaster. Journal of neurogenetics 9, 1–14 (1993).

56. Awasaki T, Lai SL, Ito K, Lee T. Organization and postembryonic development of glial cells in the adult central brain of Drosophila. The Journal of neuroscience : the official journal of the Society for Neuroscience 28, 13742–13753 (2008).

57. Ho J, Tumkaya T, Aryal S, Choi H, Claridge-Chang A. Moving beyond P values: data analysis with estimation graphics. Nature methods 16, 565–566 (2019).

58. Wickham H. The Split-Apply-Combine Strategy for Data Analysis. Journal of Statistical Software 40, 1–29 (2011).

59. Wickham H. ggplot2: Elegant Graphics for Data Analysis. Springer-Verlag New York, (2016).

60. Hirano Y, et al. Fasting launches CRTC to facilitate long-term memory formation in Drosophila. Science 339, 443–446 (2013).

61. Tully T, Quinn WG. Classical conditioning and retention in normal and mutant Drosophila melanogaster. *Journal of comparative physiology A*, Sensory, neural, and behavioral physiology 157, 263–277 (1985).

62. Awata H, Takakura M, Kimura Y, Iwata I, Masuda T, Hirano Y. The neural circuit linking mushroom body parallel circuits induces memory consolidation in Drosophila. Proceedings of the National Academy of Sciences of the United States of America 116, 16080–16085 (2019).

63. Yapici N, Cohn R, Schusterreiter C, Ruta V, Vosshall Leslie B. A Taste Circuit that Regulates Ingestion by Integrating Food and Hunger Signals. Cell 165, 715–729 (2016).

64. Münch D, Goldschmidt D, Ribeiro C. The neuronal logic of how internal states control food choice. Nature 607, 747–755 (2022).

65. Schindelin J, et al. Fiji: an open-source platform for biological-image analysis. Nature methods 9, 676–682 (2012).

66. Bolger AM, Lohse M, Usadel B. Trimmomatic: a flexible trimmer for Illumina sequence data. Bioinformatics 30, 2114–2120 (2014).

67. Dobin A, et al. STAR: ultrafast universal RNA-seq aligner. Bioinformatics 29, 15–21 (2013).

68. Anders S, Pyl PT, Huber W. HTSeq--a Python framework to work with high-throughput sequencing data. Bioinformatics 31, 166–169 (2015).

69. Team RC. A Language and Environment for Statistical Computing. R Foundation for Statistical Computing, Vienna, Austria (2023).

70. Love MI, Huber W, Anders S. Moderated estimation of fold change and dispersion for RNA-seq data with DESeq2. Genome biology 15, 550 (2014).

71. Cao K-AL, Rossouw D, Robert-Granié C, Besse P. A Sparse PLS for Variable Selection when Integrating Omics Data. Statistical Applications in Genetics and Molecular Biology 7, (2008).

72. Saldanha AJ. Java Treeview--extensible visualization of microarray data. Bioinformatics 20, 3246–3248 (2004).

